# GCNA preserves genome integrity and fertility across species

**DOI:** 10.1101/570804

**Authors:** Varsha Bhargava, Courtney D. Goldstein, Logan Russell, Lin Xu, Murtaza Ahmed, Wei Li, Amanda Casey, Kelly Servage, Rahul Kollipara, Zachary Picciarelli, Ralf Kittler, Alexander Yatsenko, Michelle Carmell, Kim Orth, James F. Amatruda, Judith L. Yanowitz, Michael Buszczak

## Abstract

The propagation of species depends on the ability of germ cells to protect their genome in the face of numerous exogenous and endogenous threats. While these cells employ a number of known repair pathways, specialized mechanisms that ensure high-fidelity replication, chromosome segregation, and repair of germ cell genomes remain incompletely understood. Here, we identify Germ Cell Nuclear Acidic Peptidase (GCNA) as a highly conserved regulator of genome stability in flies, worms, zebrafish, and humans. GCNA contains a long acidic intrinsically disordered region (IDR) and a protease-like SprT domain. In addition to chromosomal instability and replication stress, GCNA mutants accumulate DNA-protein crosslinks (DPCs). GCNA acts in parallel with a second SprT domain protein Spartan. Structural analysis reveals that while the SprT domain is needed to limit meiotic and replicative damage, most of GCNA’s function maps to its IDR. This work shows GCNA protects germ cells from various sources of damage, providing novel insights into conserved mechanisms that promote genome integrity across generations.

**Highlights:** GCNA ensures genomic stability in germ cells and early embryos across species

GCNA limits replication stress and DNA double stranded breaks

GCNA restricts DNA-Protein Crosslinks within germ cells and early embryos

The IDR and SprT domains of GCNA govern distinct aspects of genome integrity

**Graphic Abstract:** 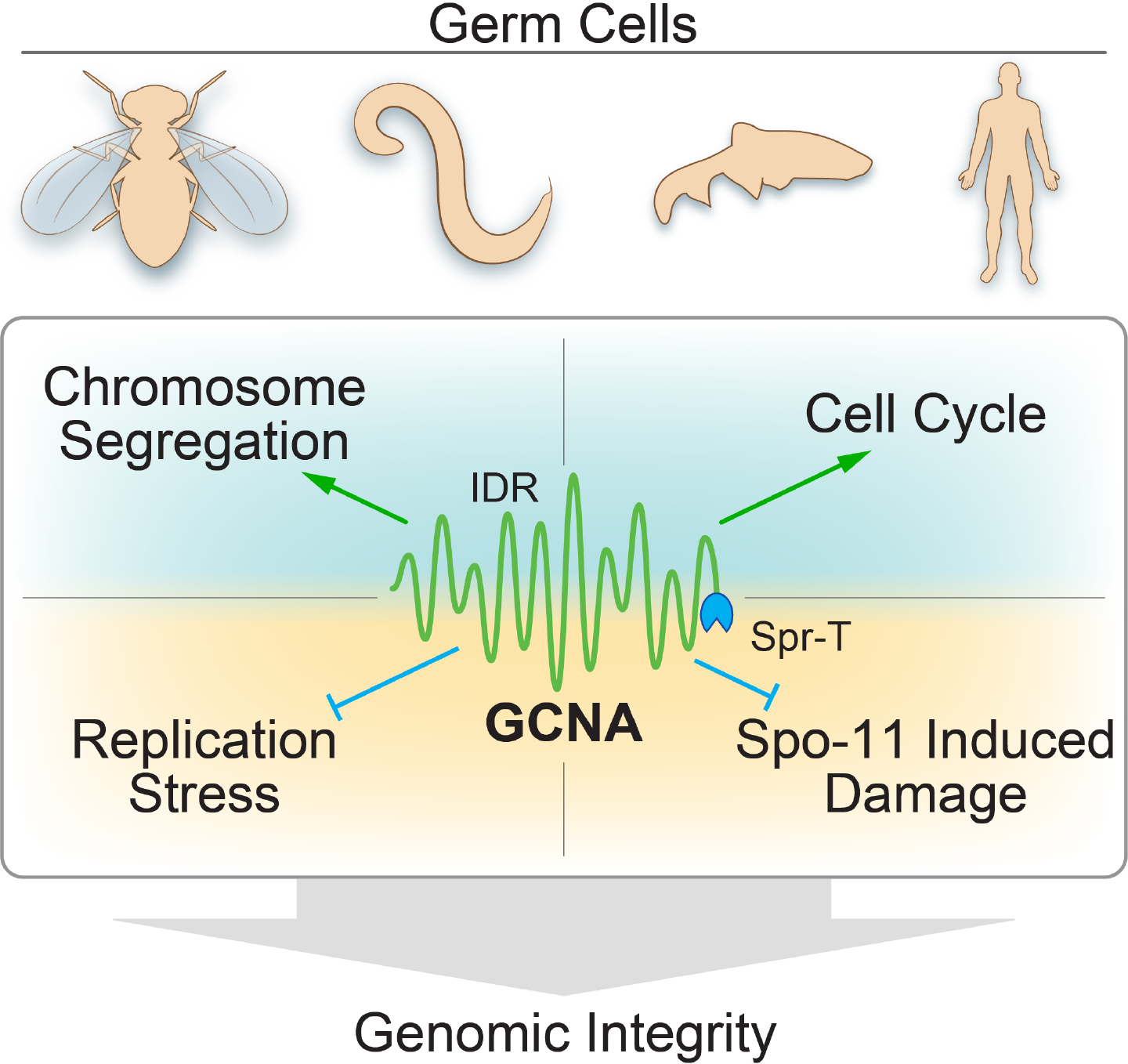

## INTRODUCTION

Early in the development of metazoans, primordial germ cells are set apart from somatic cells and undergo special programs to preserve the integrity of the genome across generations. These include producing haploid gametes through meiosis, inducing and repairing programmed DNA double-strand breaks (DSBs), inhibiting transposable elements, and reprogramming their chromatin back to an epigenetic state that supports totipotency in the fertilized zygote (Kurimoto and Saitou, 2018; Tang et al., 2016). Facilitating these processes are a subset of germ cell-specific proteins that have been conserved across millions of years, including the DEAD-box helicase Vasa, the RNA-binding protein Nanos, and the piRNA processing enzyme PIWI/Argonaute.

The recently identified Germ cell nuclear acidic peptidase (GCNA), also known as Germ cell nuclear antigen or Acidic repeat containing (ACRC), has been conserved across 1.5 billion years of evolution and has remained tightly associated with sexual reproduction, showing enriched expression within germ cells in both invertebrate and vertebrate species (Carmell et al., 2016). GCNA proteins contain an N-terminal intrinsically disordered region (IDR) which is conserved structurally despite amino acid divergence. In most species, although not in the rodent lineage, GCNA proteins also contain a C-terminal Spartan (SprT)-domain, which resembles a bacterial metalloprotease, a Zinc finger (ZnF), and an HMG box. Despite this modular domain structure and high degree of conservation, insights into GCNA function are lacking.

IDR-containing proteins have emerged as important players in cell biology, regulating phase transitions in a number of membrane-less organelles. In the nucleus, IDR proteins comprise nucleoli, speckles, and Cajal bodies. All of these condensates are thought to be macromolecular assembly sites for protein-nucleic acid complexes that control chromatin structure, transcription, and various aspects of RNA processing. IDR proteins are also found in numerous germ cell specific structures (Seydoux, 2018). For example, the IDR-containing MUT-16 protein phase separates to form mutator bodies in worms (Uebel et al., 2018). Vasa also contains an extensive disordered region which contributes to its molecular behavior and function (Nott et al., 2015). Similarly, *C. elegans* MEG-3 and MEG-4 proteins bind to and phase separate RNA to form granules both *in vitro* and *in vivo* (Smith et al., 2016). MEG-3 and MEG-4 are GCNA family members (Carmell et al., 2016), raising the possibility that GCNA itself may mediate essential germ line functions through its IDR.

Potential insight into GCNA function comes from recent investigation into the functions of Spartan proteins for which the SprT-domain gets its moniker. Several independent groups have provided evidence that Spartan proteins specifically cleave DNA-protein crosslinks (DPCs) through their SprT protease domain (Lopez-Mosqueda et al., 2016; Stingele et al., 2016; Vaz et al., 2016). DPCs represent particularly insidious lesions that can interfere with almost every chromatin-based process including replication, transcription, and chromatin remodeling (Stingele et al., 2017). The protease activity of Spartan appears highly regulated, and one major target of Spartan proteolysis is Spartan protein itself. Loss of Spartan in humans and mice results in sensitivity to UV damage, progeroid-like phenotypes, and a predisposition to hepatocellular carcinoma, suggesting the protein plays an essential role in maintaining genome integrity. Within the germ line, DPCs are formed both by Spo11 during its formation of meiotic DSBs and also by topoisomerases during mitotic and meiotic DNA replication. In addition, epigenetic reprogramming, including histone demethylation, can form potentially damaging by-products like formaldehyde (Walport et al., 2012) that can lead to the formation of DPCs (Stingele et al., 2017). Inability to remove these DPCs would interfere with the faithful transmission of the genome over generations.

Here, we provide evidence that loss of *GCNA* results in genomic instability in *Drosophila*, *C. elegans*, zebrafish, and human germ cell tumors. GCNA acts to limit Spo11 activity in flies and prevent replication stress in flies and worms. Further analysis shows that GCNA functions in parallel to Spartan proteins within germ cells. Loss of GCNA results in the accumulation of DPCs in germ cells and early embryos. Genetic and transgenic analysis points to distinct roles for the IDR and SprT domains of GCNA. Together, these results reveal a new mechanism by which germ cells ensure the integrity of their genomes from one generation to the next.

## RESULTS

### *Drosophila*, *C. elegans* and zebrafish *GCNA* mutants exhibit genome instability and chromosome segregation defects

*GCNA* exhibits enriched expression in germ cells across species (Carmell et al., 2016). We turned to *Drosophila*, *C. elegans*, and zebrafish as model systems in which to characterize the molecular function of GCNA family members. The *Drosophila* genome encodes for three potential GCNA orthologs (Figures 1A,B; S1). We generated null mutations in all three fly genes (Figure S1A-E) using CRISPR/Cas9-based techniques (Gratz et al., 2013; Gratz et al., 2015a; Gratz et al., 2015b). Only mutations in *CG14814* (hereafter called GCNA, Fig 1B), resulted in clear phenotypes, while the other two appeared viable and fertile under normal culture conditions. Homozygous *GCNA*^*KO*^ mutants were viable and fertile, but stopped laying eggs after approximately one week. The eggs that were laid exhibited maternal-effect semi-lethality (Figure S1F) marked by widespread chromosome bridges and other mitotic defects in early embryos (Figure 1C).

**Figure 1.**
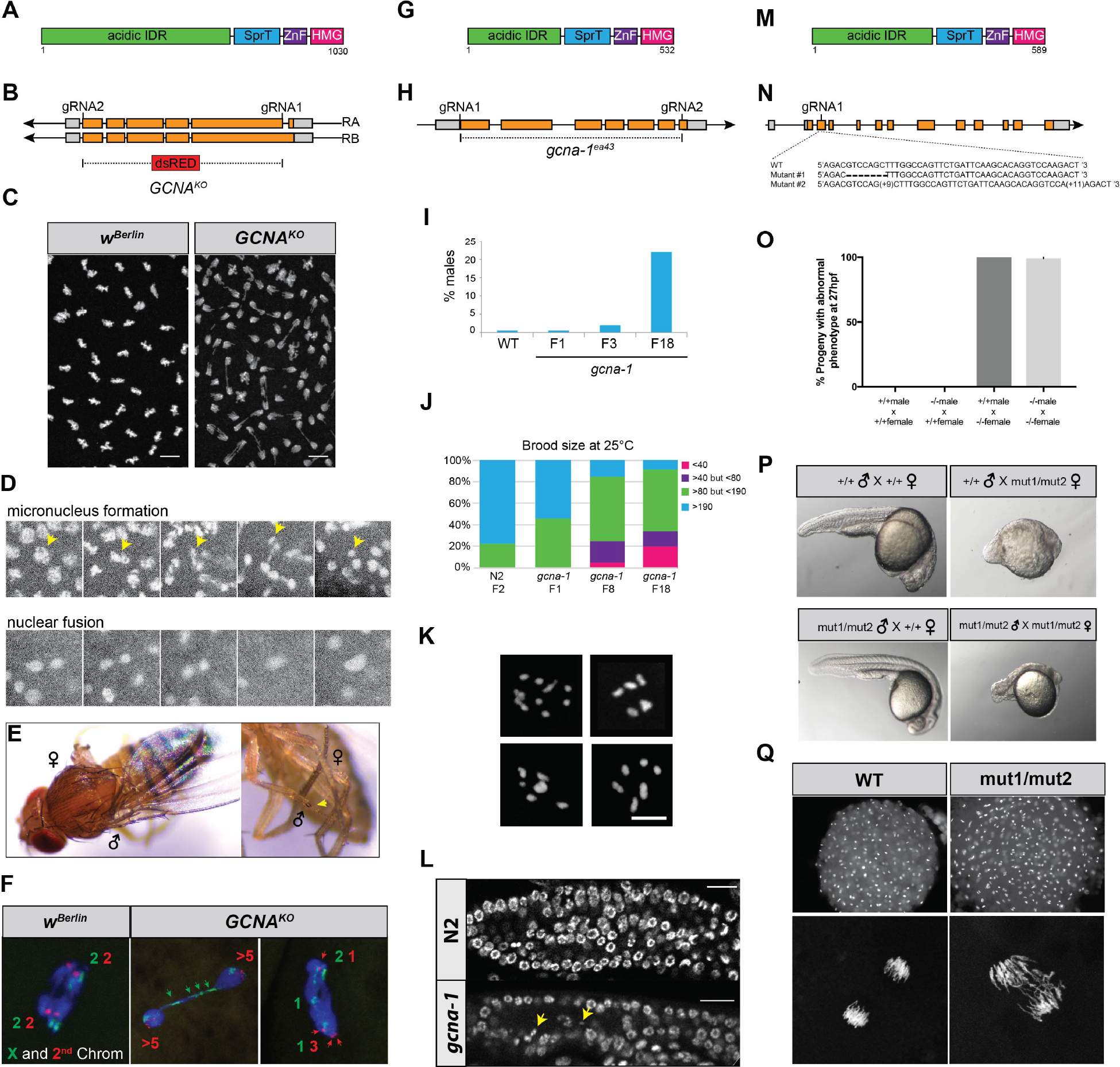
Loss of GCNA results in chromosome instability across species. **(A)** Domain organization of the *Drosophila* GCNA protein. **(B)** Organization of *Drosophila* GCNA gene locus with the design of the *GCNA*^*KO*^ allele. **(C)** DAPI staining of fixed samples reveals mitotic defects caused by loss of *Drosophila* maternal GCNA. **(D)** Still images from live cell imaging showing formation of a micronucleus (yellow arrows) and nuclear fusion events. **(E)** Examples of gynandromorphic phenotypes in progeny of *GCNA* mutant females. **(F)** FISH using a X chromosome probe (green) and second chromosome probe (red) on fly embryos derived from control and *GCNA*^*KO*^ mutant females. **(G)** Domain structure of *C. elegans* GCNA protein. **(H)** Organization of the *C. elegans gcna-1* gene detailing the design of the *gcna-1(ea43)* allele. **(I)** Quantification of worm HIM phenotype that increases over generations (n=12 for WT and n= 21, 21, 34 for *gcna-1* F1, F3, and F18 respectively) **(J)** *gcna-1* mutant worms exhibit decreases in brood size over generations (n= 12 for WT, and 34 each for *gcna-1* F1, F8, and F18). **(K)** Examples of abnormal numbers of DAPI bodies from *gcna-1* mutant germ cells from a F15 - F18 generations, prior to the onset of sterility. **(L)** *gcna-1* mutant worms exhibit chromosome fragmentation (yellow arrows). F3 dissected germ line stained with DAPI are shown. **(M)** Domain structure of the zebrafish GCNA protein. **(N)** Organization of the zebrafish *gcna* gene showing the sequence from the *mut1* and *mut2* alleles. **(O)** Quantification of maternal-effect lethality of *gcna* transheterozygous mutants **(P)** Zebrafish embryos derived from mutant *gcna* mutant females exhibit profound maternal-effect defects, regardless of the genotype of the father. **(Q)** Embryos derived from mutant *gcna* mutant females exhibit asynchronous divisions and extensive chromosome bridging during early development.

We noted the appearance of micronuclei and nuclear fragmentation in fixed *Drosophila* embryos from *GCNA* mutant females (Figure 1C). To characterize this phenotype further, we performed live cell imaging using a mRFP tagged H2Av transgene (Movie S1). This approach provided further evidence that embryos from *GCNA* mutant females exhibited a number of different defects, including the formation of micronuclei and the fusion of nuclear material (Figure 1D; S1G; Movie S1). The normally synchronous nature of the cell cycle also appeared disrupted in maternal *GCNA* mutant embryos (Movie S1).

The small number of F1 progeny from *GCNA*^*KO*^ mutant female *Drosophila* that survived to adulthood appeared sickly and sub-fertile. Of these, four percent displayed bilateral gynandromorphism (Figures 1E; S1G). This unusual phenotype, likely caused by X chromosome loss during one of the first embryonic divisions, suggested that disruption of *GCNA* results in chromosome segregation defects or chromosome instability during early embryogenesis. To test whether these defects are limited to the X-chromosome, we performed fluorescent *in situ* hybridization (FISH) on both early embryos and ovaries using probes specific for the X (359-bp repeats) and second chromosome (AACAC_(n)_ repeats). In control female embryos, two discrete X-chromosome spots and two chromosome 2 spots were observed in dividing nuclei, as expected (Figure 1F). By contrast, embryos derived from *GCNA* mutant females displayed a variety of chromosomal defects including chromosome bridges containing the X-chromosome 359-bp repeat sequences and missegregation of second chromosomes (Figure 1F). Similar phenotypes were also observed in *GCNA* mutant ovaries (Figure S1H). In addition, the nuclei of some *GCNA* mutant embryos appeared to undergo either extra rounds of replication without intervening divisions or nuclear fusion events, resulting in the appearance of four or more discrete X and chromosome 2 foci (Figure 1F). These results indicate that the chromosomal defects exhibited by *GCNA* mutants are not specific for the X chromosome. These data indicate that loss of *GCNA* results in widespread chromosomal defects, marked by problems with chromosome segregation and regulation of the cell cycle.

In parallel with the *Drosophila* experiments described above, we made a null mutation in the *C. elegans GCNA* homolog, CELE_ZK328.4 (now called *gcna-1*) (Figure 1G, H). A previous large-scale RNAi screen noted that *gcna-1* knockdown resulted in a very mild High Incidence of Males (HIM) phenotype (Colaiacovo et al., 2002), which is usually indicative of X chromosome nondisjunction (Hodgkin et al., 1979). Our CRISPR/Cas9-induced null allele (Figure 1H) confirmed this mild HIM phenotype, which is exacerbated by growth at higher temperature (25°C) and at later generations (Figure 1I). Moreover, we found that loss of *gcna-1* gives rise to a mutator phenotype that continued to worsen over generations, eventually leading to reduced lifespans, decreased mobility, and loss of fertility (Figure 1J; Figure S1I, J).

To address the nature of this phenotype, we examined worms from the generation prior to the onset of sterility using whole-mount staining. Wild-type worms contain two U-shaped germlines filled with developing oocytes that ultimately arrest at diakinesis of prophase I with 6 bivalent chromosomes. By contrast, late generation *gcna-1* mutant worms showed a range of phenotypes, from near wild-type appearing germ lines to severely runty germ lines (Figure S1K). Diakinesis nuclei of late generation *gcna-1* mutant animals also showed chromosomal abnormalities with 4 – 9, often irregularly-shaped, DAPI-stained bodies (Figure 1K), indicative of chromosome fusions and defects in crossover formation (Dernburg et al., 1998). Multiple independent lines began to produce male offspring in the several generations before the onset of sterility. In these populations, all worms assayed presented with a consistent karyotype of 5 DAPI-positive bodies suggesting they may have contained X/autosome fusion chromosomes. Similar to the fly embryo phenotypes, we observed the presence of chromosome bridges and DNA fragments in *gcna-1* mutant germ cells (Figure 1L, S1L). Together, the fly and worm mutant phenotypes indicate that loss of *GCNA* function disrupts reproductive success and chromosome stability across species.

To characterize *GCNA* function in a vertebrate species, we generated a number of *gcna* mutant alleles in zebrafish, using CRISPR/Cas9 techniques. We used a single gRNA to induce double-strand breaks within the coding sequence of the third exon of the zebrafish *gcna* gene and screened for insertions and deletions within the locus. These efforts resulted in the isolation of a number of alleles, including a 7 bp deletion (mut1) and a complex insertion of 9 bp and 11 bp (mut2) (Figure 1M, N). Both resulted in frameshifts predicted to create truncations early in the protein coding sequence. Crosses between these two alleles resulted in viable transheterozygotes. However, the progeny of female transheterozygotes displayed widespread morphological defects and cell death (100% penetrant; n>20 embryos) (Figure 1O, P; Figure S1M, N). Close examination of early embryos from mutant females revealed many cells that had undergone asynchronous mitotic divisions and contained tangled chromosomes, in contrast to wild-type controls (Figure 1Q). These results indicate that disruption of zebrafish *gcna* results in maternal-effect lethality marked by chromosome instability remarkably similar to the phenotypes observed in flies and worms.

### *Drosophila GCNA* mutant ovaries exhibit increased DNA damage

As suggested by the female sterility that ensues after one week, loss of GCNA resulted in a number of ovarian phenotypes in *Drosophila* (Figure 2A; S2A-D). For example, many *GCNA* mutant egg chambers deviated from the normally invariant number of 16 germ cells per developing cyst. In aged flies, this phenotype grew more penetrant. Labeling for ring canals and the cell death marker cleaved caspase 3 suggested that these counting defects arose from abnormal cell divisions and the loss of the normal synchrony of germ cell divisions, consistent with the cell cycle defects observed in maternal mutant embryos (Figure S2B, data not shown). We also observed defects in differentiation and delays in oocyte specification (Figure S2C, D).

**Figure 2.**
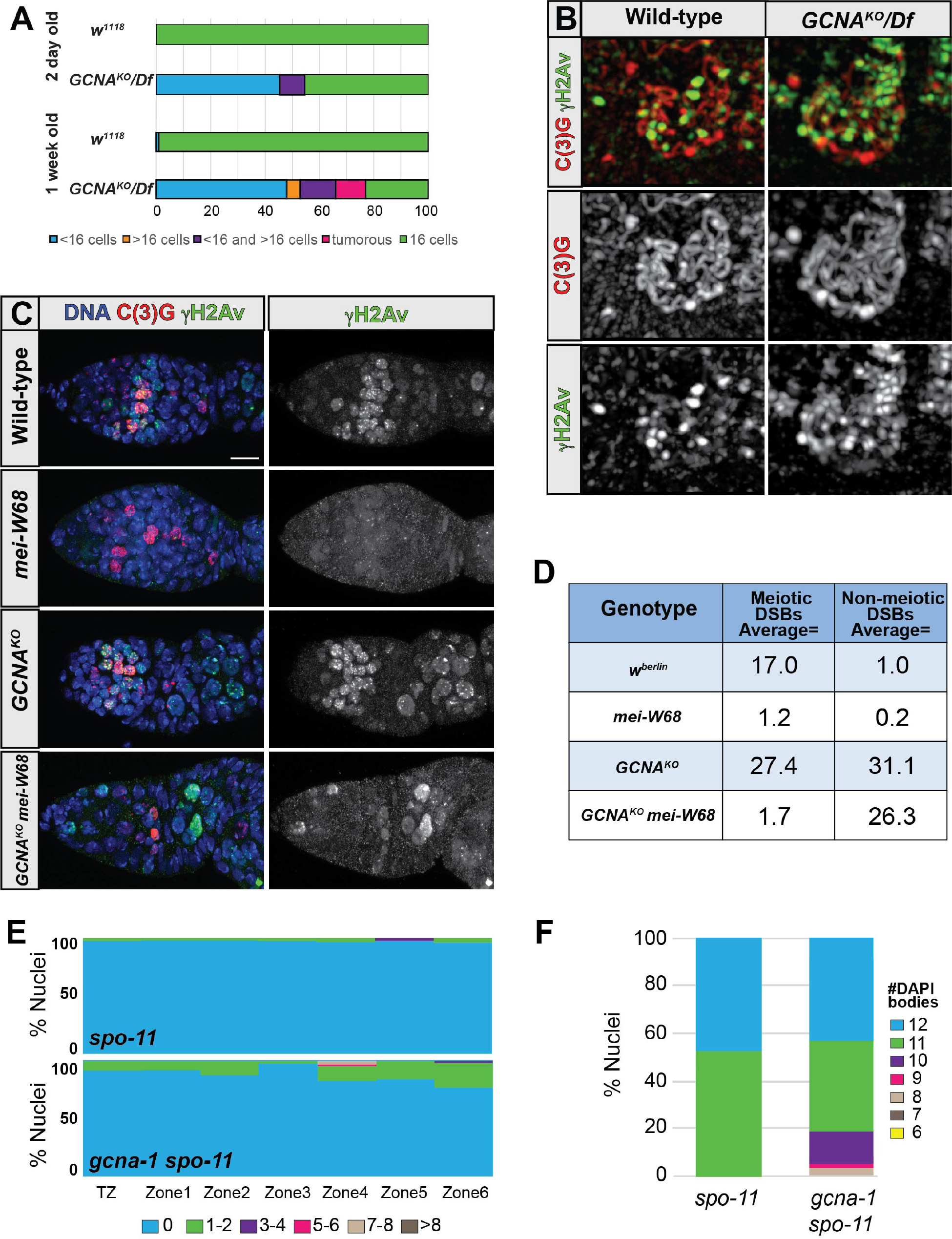
GCNA mutants exhibit excessive Spo11-dependent and independent breaks during oogenesis. **(A)** Quantification of *Drosophila* ovarian phenotypes, which include abnormal numbers of cells within germline cysts and tumor formation. **(B)** SIM images of Wildtype and *GCNA* mutant meiotic nuclei stained for C(3)G (red) and gH2Av (green). **(C)** Wild-type, *GCNA*^*KO*^, *mei-W86*^*0949*^, and *GCNA*^*KO*^; *mei-W68*^*0949*^ stained for C(3)G (red), γH2Av (green) and DNA (blue). **(D)** Table of the average number of γH2Av labeled DSBs per nucleus in the indicated genotypes. **(E)** Quantification of meiotic (leptotene through onset of diplotene) RAD-51 foci in *spo-11* and *gcna-1;spo-11* mutant *C. elegans* germ lines (n = 3 germ lines each). **(F)** Quantification of DAPI positive bodies in diakinesis-stage, −1 oocytes of *C. elegans* (*spo-11*, n= 20; *gcna-1;spo-11*, n=42).

To test whether loss of *GCNA* also disrupts meiosis in *Drosophila*, we stained control and *GCNA* mutant *Drosophila* ovaries with antibodies to detect the synaptonemal complex (SC) protein C(3)G and the DSB marker γH2Av, which is analogous to γH2AX in other systems (Jang et al., 2003; Page and Hawley, 2001). In control ovaries, the SC and DSBs were first observed in region 2A of the germarium, as previously described (Jang et al., 2003). In *GCNA* mutant germaria, the SC formed normally in region 2A. Strikingly, however, *GCNA* mutant germ cells exhibited larger and many more γH2Av positive foci than controls. Structured Illumination Microscopy (SIM) and confocal microscopy revealed that many of the large γH2Av foci in *GCNA* mutant cells can be resolved into individual foci (Figure 2B, C). Quantification of these foci showed that individual GCNA mutant nuclei experienced, on average, many more breaks than controls (Figure 2D). We observed individual nuclei with over 40 γH2Av foci.

In control germaria, DSBs are rapidly repaired. However, in *GCNA* mutant germ cells, γH2Av staining extended into early egg chambers. In addition, the expression of a *p53* reporter, which correlates with DNA damage (Lu et al., 2010; Wylie et al., 2014), also exhibited expanded expression into early egg chambers in *GCNA*^*KO*^ ovaries (Figure S2E). We considered that the greater number and persistence of DSBs within *Drosophila GCNA* mutant cells could reflect either a failure to repair breaks or a failure to limit the number of breaks that normally form, or both. Mutations that disrupt the homologous recombination repair pathway and cause activation of the Chk2 checkpoint lead to a number of shared phenotypes including egg chamber patterning defects, as reflected by the fusion or absence of dorsal appendages in mature eggs, meiotic non-disjunction, and sensitivity to DNA damaging agents (Abdu et al., 2003; Ghabrial et al., 1998; Staeva-Vieira et al., 2003). Unlike *Rad51* and *Rad54* mutants, which exhibit dorsal appendage defects in 50-60% of their eggs, only 4-8% of *GCNA* mutant eggs exhibit this phenotype (Figure S2F; n=>200 individual eggs from multiple egg lays). *GCNA* mutants flies also do not display whole body sensitivity to irradiation (IR, Figure S2H) unlike *Rad51* and *Rad54* mutants (Ghabrial et al., 1998; Staeva-Vieira et al., 2003). Together with our observation that meiotic nondisjunction rates did not vary between controls and *GCNA*^*KO*^ homozygotes (Figure S2I), these results suggest that GCNA likely does not play an essential role in meiotic HR-mediated repair in *Drosophila*.

We next considered the possibility that loss of GCNA disrupts the ability to limit DSBs. Enhanced activity of Spo11 during meiosis or a failure to spatially and temporally restrict Spo11 activity could be responsible for the formation of the additional DSBs we observe in *GCNA* mutants. If correct, this model would predict that loss of *Spo11*, named *mei-W68* in *Drosophila* (McKim and Hayashi-Hagihara, 1998), would suppress *GCNA* mutant phenotypes. To test for this possibility, we made double mutants between *GCNA* and *mei-W68* and *mei-P22*, a gene needed for targeting mei-W68 to break sites (Liu et al., 2002). Loss of either *mei-W68* or *mei-P22* resulted in a dramatic suppression of the extra meiotic DSBs that form in region 2A of *GCNA-*mutant germaria (Figures 2C,D; S2J), supporting the interpretation that GCNA may either limit Spo11 accessibility to DNA or temporally restrict Spo11 activity. In these double mutants, however, we still observed γH2Av staining in early and later germ cells suggesting a subset of breaks occurs independently of the meiotic program. In addition, neither *mei-P22* nor *mei-W68* mutations suppressed other phenotypes associated with loss of *GCNA*, including the germ-cell counting defects and maternal-effect semi-lethality.

Excessive DNA damage during meiosis was not obvious in *C. elegans gcna-1* mutants (Figure S2K). By crossing *gcna-1* into a *spo-11* mutant background and exposing worms to the DSB-inducing agent gamma irradiation, we were able to see similar accumulation of RAD-51 post-exposure (Figure S2L), suggesting that early steps in DNA repair are normal in *gcna-1* mutant animals, as they are in *Drosophila GCNA* mutants. However, we noticed that RAD-51 foci were present in the pachytene nuclei of the unirradiated *gcna-1;spo-11* mutant controls (Figure 2E). We infer that these foci arose as the consequence of DNA damage incurred during either mitotic divisions of germ cells or meiotic S phase. This “carry through” damage can induce meiotic crossover (CO) formation as seen by a decrease in univalent chromosomes in *gcna-1;spo-11* double mutant worms compared to *spo-11* mutant worms (Figure 2F). Together with the *Drosophila* experiments, these findings indicate that while GCNA plays species-specific roles in the regulation of Spo11 activity during meiosis, loss of *GCNA* function leads to the accumulation of Spo11-independent DSBs within both *Drosophila* and *C. elegans* germ cells.

### Loss of GCNA leads to replication stress in *Drosophila* and *C. elegans*

To investigate the nature of the SPO11-independent DNA damage and the source of the transgenerational sterility phenotypes, we carried out a number of experiments to determine if GCNA directly disrupted homologous recombination-mediated repair or transposable element (TE) surveillance (Figure S3). We saw no increased sensitivity to gamma irradiation in worms (Figure S3A) suggesting GCNA does not a major role in HR-mediated DNA repair. We saw a modest increase in TE expression, which suggested a potential role for GCNA in the piRNA/PIWI-mediate TE surveillance pathway (Figure S3C). However, *gcna-1* mutations were synthetic sterile with the *prg-1/*PIWI mutations (Figure S3D), arguing that GCNA acts in parallel to the canonical pathway for TE surveillance. In flies, expression and localization of Aubergine, the major argonaute in the piRNA pathway, was unaffected in *GCNA* mutants (Figure S3E). Transcriptomic analyses identified an approximate 2-fold increase in expression of telomere-associated TEs and a concomitant decrease in metabolic genes (Figure SF-J), raising the possibility that cellular stress pathways may be activated in GCNA mutants.

We therefore considered the possibility that *gcna-1* mutants experience an increase in replicative stress, which is both a major cause of endogenous DSBs and also causes chromosome segregation defects and disruption of the cell cycle (Magdalou et al., 2014; Zhang et al., 2018). We therefore sought to determine whether loss of GCNA function contributes to replicative repair. We took advantage of a mutation in the DEAD-box helicase *dog-1/FANCJ* which is the only known gene required for replication of G-quadraplex-like structures in the worm (Kruisselbrink et al., 2008). Accordingly, DOG-1is required for accurate replication through G-C rich DNA (Youds et al., 2008) and *dog-1* mutations exhibit microsatellite repeat instability (MSI) seen by changes in repeat length in a PCR-based assay of repeats in the *vab-1* gene (Youds et al., 2008); Figure S4A). Whereas *C. elegans gcna-1* mutation did not exhibit repeat instability on its own, it strongly enhanced the *dog-1* phenotype (Figure 3A). This result is consistent with a role for GCNA in promoting replicative repair and/or replication restart.

**Figure 3.**
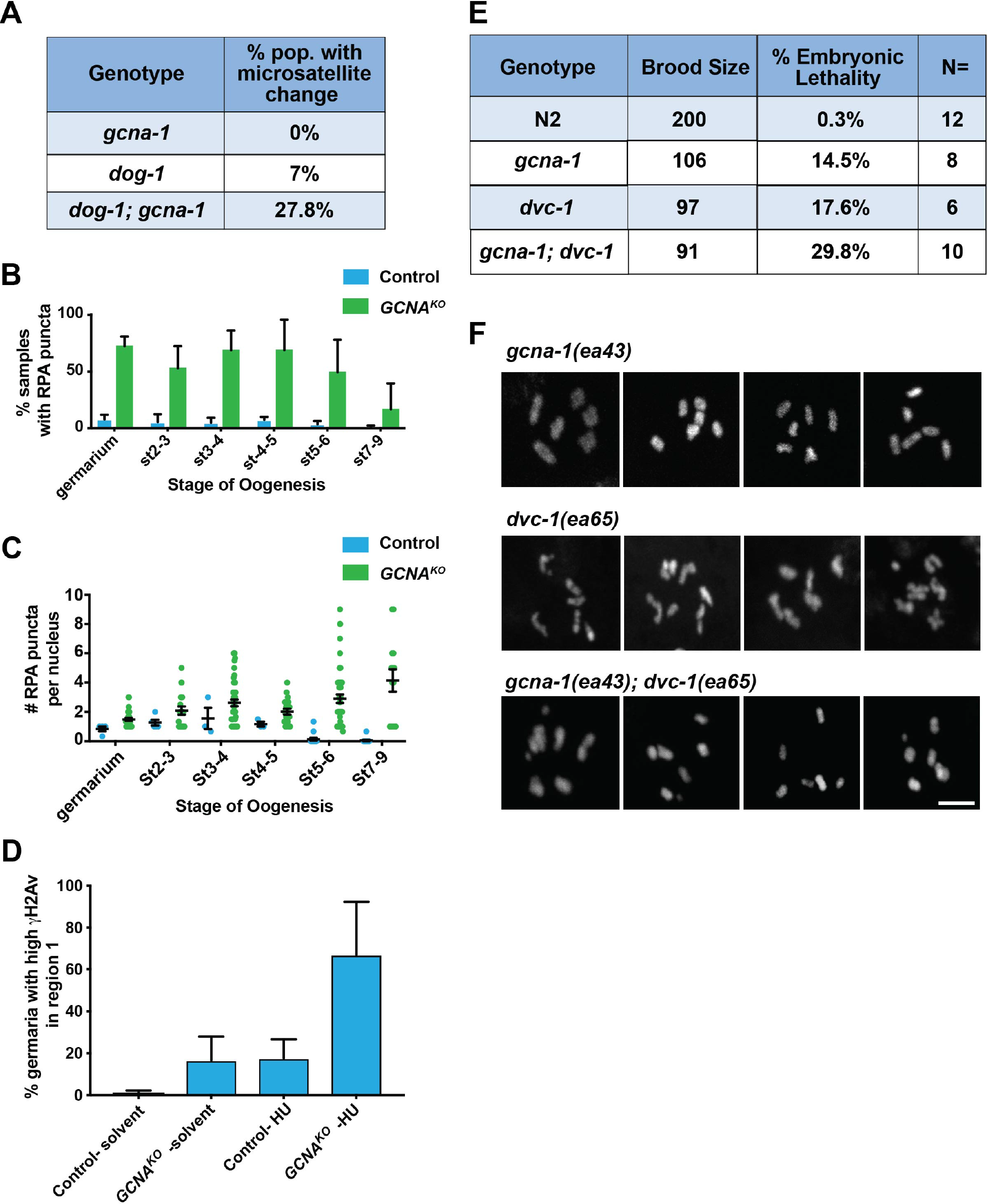
GCNA acts to prevent replicative stress. **(A)** Analysis of microsatellite repeat changes within the *vab-1* locus of *C. elegans*. Percentage of the *dog-1* (n=162), *gcna-1* (n= 76), or *dog-1;gcna-1* (n=189) worms that exhibits changes in microsatellite length. **(B)** Quantification of the number of RPA punctae per nucleus in the indicated stages of oogenesis. **(C)** Quantification of the percent of nuclei with RPA punctae in the indicated stages of oogenesis. **(D)** Percentage of germaria (n=>30/ replicate; 3 independent biological replicates) from control and *GCNA*^*KO*^ homozygous females fed solvent or 50 mM HU that displayed γH2AV in region 1. **(E)** Brood analysis (total viable adult offspring) and embryonic lethality associated with N2, *dvc-1*, and *gcna-1* single mutants and *dvc-1;gcna-1* double mutant worms. **(F)** Examples of DAPI-stained diakinesis oocytes from *dvc-1*, *gcna-1*, and *dvc-1; gcna-1* F3 mutant germ cells.

When replication forks stall at DNA lesions or aberrant DNA structures, increased stretches of single-stranded DNA (ssDNA) that can be visualized as RPA positive punctae within nuclei are formed. To determine whether loss of *GCNA* also leads to replicative stress in *Drosophila* germ cells, we compared RPA expression and localization within control and *GCNA* mutant ovaries. Disruption of *GCNA* function resulted in the appearance of discrete RPA nuclear foci that were rarely observed in control cells (Figure 3B, C; S4B). Within *GCNA* mutant ovaries, these punctae were present in both mitotic germ cells within germaria and in endocycling nurse cells.

Next, we tested the extent to which *Drosophila GCNA* mutant germ cells were more sensitive to hydroxyurea (HU) treatment relative to controls. HU effectively limits the pool of dNTPs available for replication, causing replication fork stalling and DNA damage, particularly in mutants that already suffer from replication stress. We fed control and *GCNA* mutant females HU for 24 hours and then assayed for DNA DSBs within their ovaries after another 24 hours. *GCNA* mutant germaria accumulated many more DSBs upon HU treatment within mitotically active cells relative to controls, indicating that loss of GCNA makes germ cells more sensitive to replicative stress (Figure 3D). In the associated manuscript, the authors found the *gcna-1* mutant worms were also mildly sensitive to HU. Taken together, these results support a potential role for *GCNA* in promoting replicative repair across species.

### GCNA and Spartan act independently

DNA-protein crosslinks (DPCs) are a significant source of replicative stress (Vaz et al., 2017). GCNA belongs to the SprT domain protein family that gets its name from Spartan. Spartan has recently been shown to clear DPCs (Stingele et al., 2016; Vaz et al., 2016) To determine whether GCNA has a similar function and acts together with Spartan, we crossed our *C. elegans* and *Drosophila* null *GCNA* mutations into *Spartan* mutant backgrounds. In worms, Spartan is encoded by *dvc-1* (DNA damage-associated VCP/p97 Cofactor homolog). The previously described allele is a partial truncation that functions as a mutator (Stingele et al., 2016) and the homozygous *dvc-1(ok260)* allele we acquired manifested an Uncoordinated (Unc) phenotype. To avoid confounding results from *de novo* mutations in this genetic background, we used CRISPR/Cas9 to generate a null allele, *dvc-1(ea65)* (Figure S4C), and immediately analyzed the stock in order to avoid mutation accumulation. Homozygous *dvc-1(ea65)* mutant animals are not Unc, but do show the reduced brood sizes (Figure 3E) described for *dvc-1(ok260)*. Unlike *gcna-1*, the *dvc-1* null mutation is slow growing and difficult to maintain even at 20°C, suggesting a much more severe long-term impact on population. We crossed *dvc-1(ea65)* into *gcna-1(ea43)* and analyzed both fertility and the phenotype of diakinesis oocytes. *dvc-1* and *gcna-1* single mutants both show reduced brood sizes and increased embryonic lethality that are further exascerbated in the double mutant (Figure 3E; average of greater than 6 broods/ genotype). Of note, *gcna-1;dvc-1* showed an additive effect, almost doubling embryonic lethality (3-way Kruskal-Wallis Test, p<0.05). Similarly, diakinesis nuclei of *dvc-1(ea65)* F3 homozygous animals contain aberrant chromosome numbers and morphologies that were further exacerbated by loss of *gcna-1.* In *dvc-1* single mutants, chromosome fusions and fragments were seen in 33% and 14% of nuclei, respectively (n= 21; note that some nuclei have both fusions and fragments). In the *gcna-1;dvc-1* double mutants, the frequency of fusions was not statistically different (n= 26; Chi-square, p>0.1). However, DNA fragments were much more common in *gcna-1; dvc-1* double mutants than *dvc-1* single mutants (62% of nuclei; Chi-Square, p < 0.001; Figure 3F). These results suggest that GCNA-1 acts independently of the DVC-1 pathway to ensure genomic stability.

The *Drosophila* Spartan homolog, *maternal haploid* (*mh*), has been shown to play a role in maintaining paternal chromosome integrity during early embryogenesis. We compared egg laying and embryonic development between *GCNA* and *mh* single mutants and the *GCNA mh* double mutants. The number of eggs laid per female appeared comparable for single and double mutants over the first two days post-eclosion. Of these embryos, 11% from *GCNA* mutant females (n=299) and 1.4% from *mh^1^* mutant females (n=444) hatched, while none of the embryos from *GCNA mh* double mutants (n=296) completed embryogenesis. Next, we compared the amount of DNA damage in *GCNA*, *mh*, and *GCNA mh* double mutant ovaries by staining for γH2Av and RPA. While *mh* mutant ovaries looked comparable to control ovaries, ovaries from *GCNA mh* double mutants exhibited increased γH2Av and RPA foci relative to both controls and the single mutants (Figure S4D, E). These results suggest that GCNA and Mh likely act in parallel to limit DNA damage in the *Drosophila* germline, consistent with our worm data that Spartan and GCNA act independently. These data raise the possibility that GCNA may have an independent role in clearing a subset of DPCs.

### Loss of GCNA results in an accumulation of DPCs

We used the rapid approach to DNA adduct recovery (RADAR) coupled with SDS-PAGE and silver staining to evaluate whether GCNA influences DPC levels within *Drosophila* ovaries and early embryos (Figure 4A) (Kiianitsa and Maizels, 2013). In RADAR, cells are lysed under denaturing conditions. DNA and proteins crosslinked to DNA will precipitate in the presence of ethanol, while free proteins will remain in the supernatant. The samples are then normalized to the amount of DNA in the sample. Using this technique, we observed a modest, but consistently elevated level of DPCs in the *GCNA* mutant ovaries and early embryos derived from *GCNA* mutant females compared to control samples (Figure 4B). To test whether loss of *GCNA* resulted in elevated DPC levels across species, we performed RADAR on zebrafish embryos. We focused on zebrafish because of the acute maternal effects we observed in the *gcna* mutants and the ease with which we could obtain the necessary biological material. Again, this analysis showed that disruption of *GCNA* resulted in modestly increased levels of DPCs (Figure 4C).

**Figure 4.**
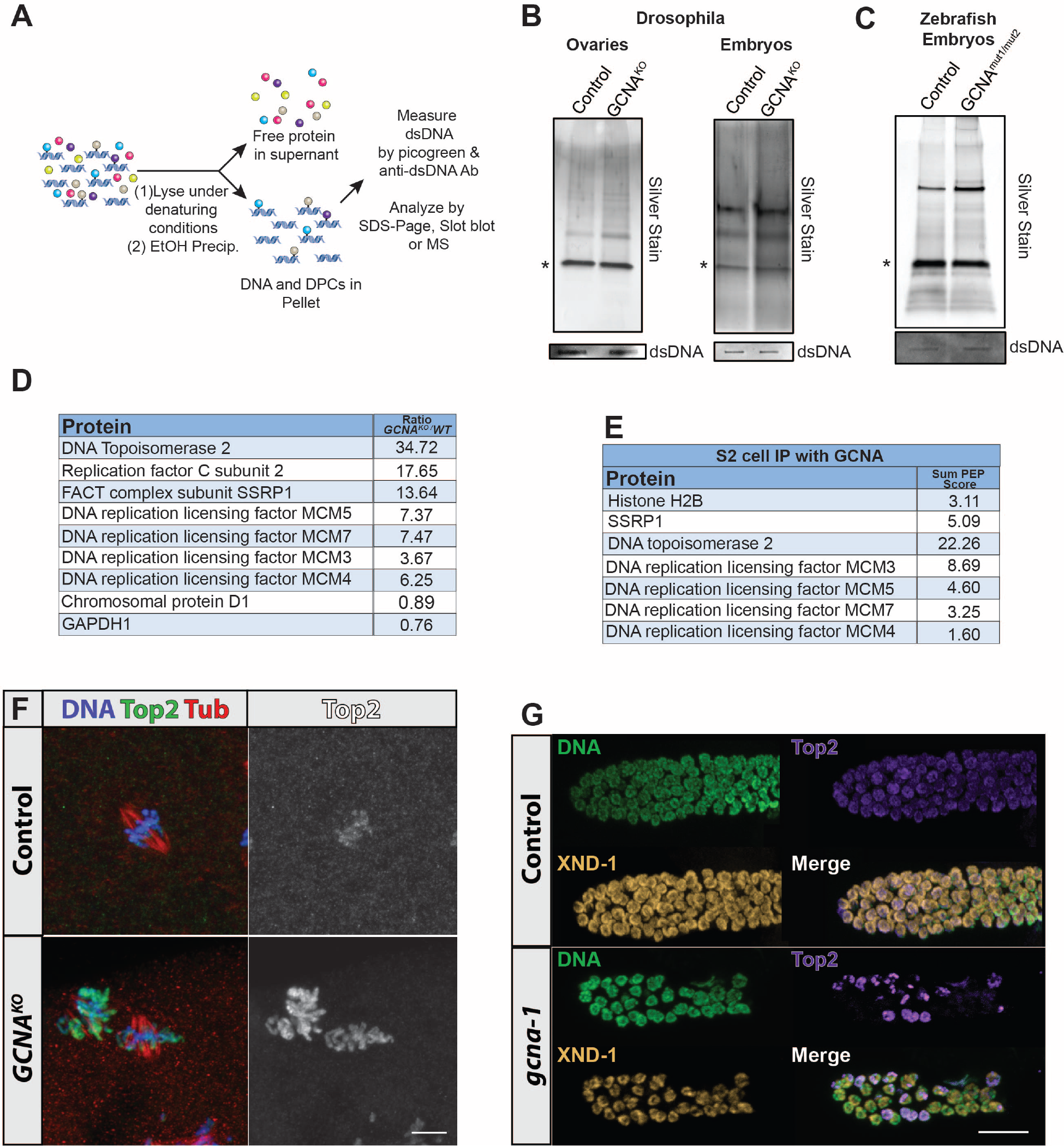
GCNA mutants exhibit increased DPC levels. **(A)** Schematic illustrating principle of the RADAR assay. **(B)** RADAR was performed on *Drosophila* ovaries and embryos. The resulting sample were normalized to dsDNA, separated by SDS-PAGE and silver stained. Asterisks mark benzonase. **(C)** RADAR was performed on zebrafish embryos at the 1000 cell stage. The resulting sample were normalized to dsDNA, separated by SDS-PAGE and silver stained. Asterisks mark benzonase. **(D)** Specific proteins were enriched in DPCs from isolated nuclei of *GCNA*^*KO*^ *mutant* embryos based on MS. **(E)** List of proteins that are pulled down in GCNA immunoprecipitation from S2 cells. **(F)** Early *Drosophila* embryos derived from control and *GCNA*^*KO*^ females stained for DNA (blue), Top2 (green) and Tub (red). Grayscale shows Top2 alone. **(G)** Control (N2) and *gcna-1* mutant *C. elegans* gonads stained for DNA (green), TOP-2 (magenta), and staining control XND-1 (yellow).

To identify proteins that formed DPCs in the absence of *GCNA*, we analyzed *Drosophila* ovary and embryo RADAR samples using mass-spectrometry (MS) (Figure S5A). A number of proteins formed DPCs specifically in *GCNA* mutant embryos or ovaries, or both; these include Histone H2B, Topoisomerase 2 (Top2), SSRP1, MCM3 and Fibrillarin. These proteins were not detected in the wild-type samples although other proteins were, pointing to the specificity of the described interactions. We repeated the RADAR assay on isolated nuclei from *Drosophila* embryos and found that we again detected an enrichment of specific DPCs in the *GCNA* mutant samples (Figure 4D). The increased levels of Top2 and MCM protein DPCs in *GCNA* mutant samples drew our attention. Topoisomerase 2 decatenates supercoiled DNA, while the MCM complex unwinds DNA in front of the replication fork. To begin to test whether functional interactions exist between GCNA and the specific proteins identified in our mass-spec analysis of DPC samples, we immunoprecipitated a tagged form of GCNA from S2 cells and performed mass-spectrometry on the resulting pellet (Figure 4E). Interestingly, several, but not all, proteins found in DPCs in *GCNA* mutants also physically interact with GCNA protein. For example, we detected Top2 in GCNA IPs, as as multiple components of the MCM complex. These results further link GCNA with direct regulation of replication in germ cells and early embryos.

To characterize how loss of *GCNA* affects these interacting proteins, we focused on Top2 expression and localization in embryos and ovaries from control and *GCNA* mutant females by immunofluorescence (Figure 4F), given the availability of appropriate reagents. In control embryos, Top2 localized to chromosomes during metaphase, as previously reported (Tang et al., 2017). Low levels could also be observed in the cytoplasm of early embryos. By contrast, embryos from *GCNA* mutant females displayed much higher levels of Top2 associated with chromosomes in metaphase and a corresponding decrease in cytoplasmic localization, suggesting Top2 protein was redistributed in the absence of *GCNA*. *GCNA* mutant egg chambers also exhibited an increase of nuclear Top2 punctae (Figure S5B-D). The overall levels of Top2 did not appear to change in *GCNA* mutants (Figure S5E). Finally, similar increases in chromosome-associated Top2 were observed in worm *gcna-1* mutant germ cells, indicating that loss of *GCNA* results in an increased accumulation of nuclear Top2 across species (Figure 4G). Thus, we have demonstrated that GCNA both interacts with and impacts the localization of proteins that become associated with DPCs when *GCNA* function is impaired.

### Characterization of *Drosophila* and *C. elegans* GCNA domain function

The accumulation of DPCs in *GCNA* mutants led us to explore the functional contribution of the conserved metallopeptidase zinc-binding SprT signature and the IDR motifs to GCNA function. To this end, we first made a C-terminally tagged GFP *Drosophila* transgene (Figure 5A). This protein predominantly localized to cytoplasmic punctae, although we could detect discrete punctae within nuclei as well. Some foci appeared enriched around the nuclear envelop, but were clearly distinct from the nuage, based on lack of co-localization with Aubergine. During mitosis, GFP-tagged GCNA associated with dividing chromosomes (Figure 5B). We also tagged the endogenous GCNA gene at the C-terminus using a triple tag, comprised of CyOFP, HA, and FLAG (Figure S6A, B). This endogenously tagged protein also exhibited predominantly cytoplasmic localization and exhibited association with chromosomes during mitosis in ovaries and early embryos.

**Figure 5.**
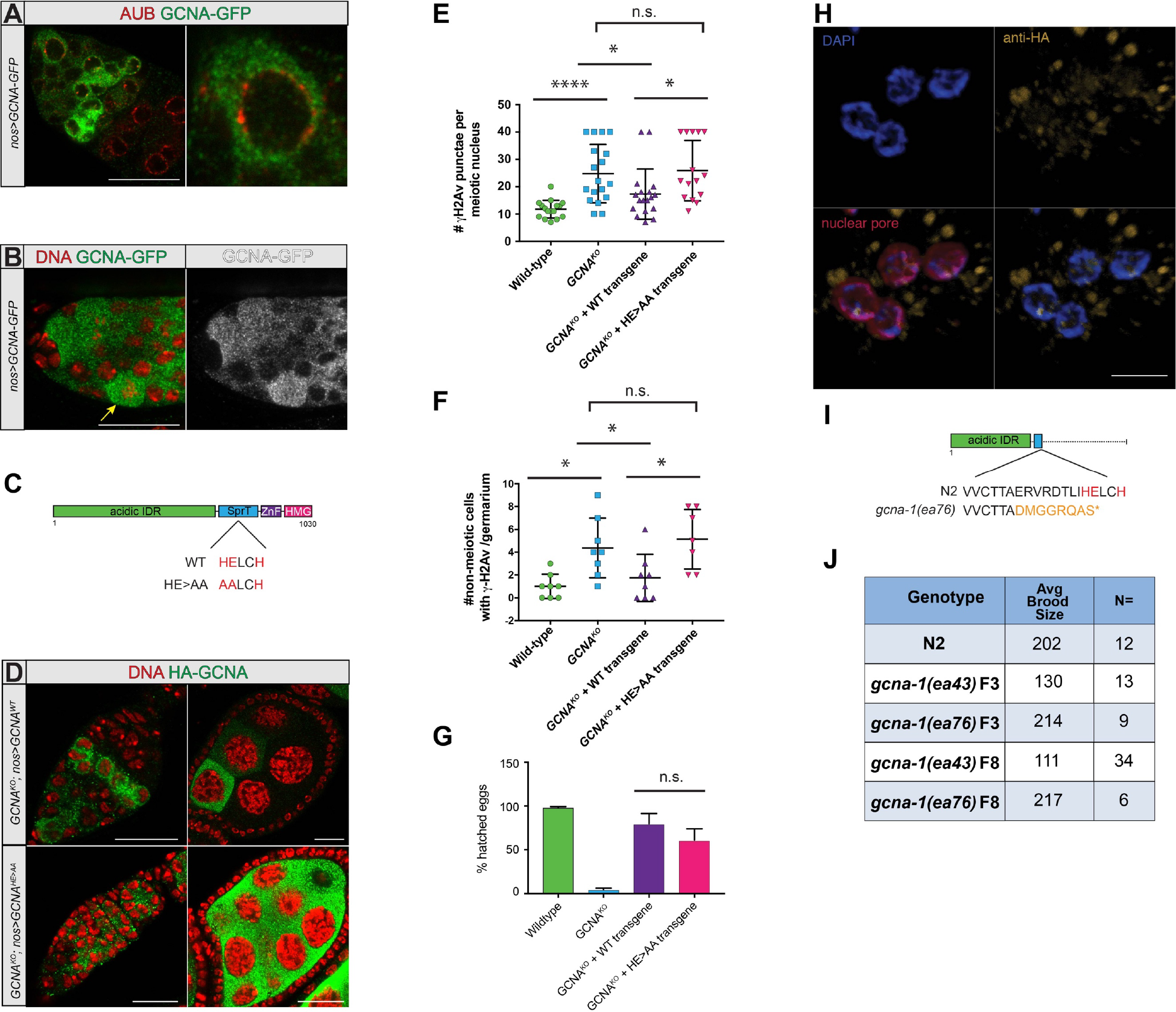
Functional analysis of GCNA domain structure. **(A)** Germarium expressing GCNA::GFP driven by *nanos (nos)-gal4* stained for GFP (green) and Aub (red). Scale bar represents 20µm. **(B)** Germarium expressing GCNA::GFP driven by *nanos (nos)-gal4* stained for GFP (green) and DNA (red). Yellow arrow points to cell undergoing mitosis. Scale bar represents 20µm. **(C)** Structure of the Drosophila *GCNA*^*WT*^ and *GCNA*^*HE>AA*^ mutant transgenes **(D)** Expression of the HA-tagged *UAS GCNA*^*WT*^ and *GCNA*^*HE>AA*^ transgenes driven by *nanos* (*nos*)-*gal4.* Scale bars represent 20µm. **(F)** The number of gH2Av foci in meiotic nuclei from wild-type, *GCNA*^*KO*^, *GCNA*^*KO*^; *nos>GCNA*^*WT*^, and *GCNA*^*KO*^; *nos>GCNA*^*HE>AA*^ ovaries **(G)** The number of γH2Av foci in non-meiotic nuclei from wild-type, *GCNA*^*KO*^, *GCNA*^*KO*^; *nos>GCNA*^*WT*^, and *GCNA*^*KO*^; *nos>GCNA*^*HE>AA*^ ovaries **(G)** Quantification of percentage of eggs derived from females of the indicated genotypes that hatch into larval development. **(H)** Expression of *C. elegans* HA::GCNA-1 from *gcna-1(*ea72 [*HA*::*gcna-1*]). Germ cells are labeled for DNA (blue), GCNA-1 (anti-HA, yellow), and the nuclear pore (mAb414, magenta). Scale bar represents 5µm. **(I)** Structure of the *C. elegans* C-terminal truncation allele, *gcna-1(ea76)*. **(J)** Comparison of brood sizes from N2 control, and homozygous, F3 and F8 *gcna-1(ea43)* null and *gcna-1(ea76)* truncation mutants.

To test the functionality of the SprT domain in *Drosophila* GCNA, we made full length UAS-HA N-terminally tagged wild-type and mutant transgenes in which we mutated key residues needed for its enzymatic activity (HE>AA), based on the characterization of Spartan proteins (Figure 5C) (Morocz et al., 2017). These transgenes were inserted into the same genomic position and exhibited similar expression levels, as assayed by western blot (Figure S6C). Like GFP and triple-tagged GCNA, these transgenes predominatly localized to specific, cytoplasmic punctae, although a small number of nuclear foci were consistently observed as well. When we expressed both transgenes in a *GCNA* mutant background using the same *nanos-gal4* driver, the HE>AA protein localized to slightly larger cytoplasmic punctae and was more broadly expressed in late stage nurse cells (Figure 5D). Together, the western blot and immunofluorescence experiments suggest that the SprT domain may regulate the size and number of GCNA labeled condensates.

Next, we tested for the ability of the *Drosophila* HA-tagged transgenes to suppress meiotic dependent and independent DNA damage and rescue the maternal-effect semi-lethality of the *GCNA* mutant. The full-length wild-type transgene was able to suppress the excessive formation of both Spo11-dependent and independent breaks that we observed in *GCNA*^*KO*^ mutant germ cells (Figure 5E, F). Strikingly, the HE>AA transgene did not rescue these phenotypes to any appreciable degree. This transgene is functional, however, as both it and the wild-type transgene were able to rescue the maternal-effect semi-lethality of embryos derived from *GCNA* mutant females (Figure 5G). For both transgenes, rescue of the maternal-effect semi-lethality was accompanied by a decrease in chromosome bridging and other mitotic defects based on labeling. Thus, the HE>AA mutation has created a separation-of-function mutation, revealing a requirement for the SprT domain for DNA damage prevention in the fly germ line, but not for chromosomal stability during early embryogenesis.

We tagged the endogenous *C. elegans gcna-1* locus at the 5’ end with either OLLAS (OmpF Linker and mouse Langerin fusion Sequence) or HA tags. The majority of OLLAS::GCNA-1 and HA::GCNA-1 proteins localized to the cytoplasm under steady-state conditions (Figure 5H; S6D). GCNA-1 proteins are maternally-inherited and are ultimately enriched in the primordial germ cell precursors, Z2 and Z3 (Figure S6E) which are readily identified by their size and position. In adult germ cells, chromosome squashes allowed us to see small puncta associated with chromosomes in the nucleus (Figure 5H; S6F), similar to what we observe in *Drosophila*.

We also made a mutant allele, *gcna-1(ea76)* which truncates the C-terminus (of the wild-type protein to retain just the IDR region (Figure 5I). While the most robust phenotype observed with the null allele, *gcna-1(ea43)*, was reduced fecundity that began at the F3 generation and became more severe in later generations (Figure 1J; 5J), *gcna-1(ea76)* had brood sizes greater than or equal to wild type in the F3 and F8 generations. This increase in brood size continued for >15 generations although a subset of the population started to exhibit phenotypes associated with *gcna-1* loss, including a HIM phenotype and sterility. Thus, while loss of catalytic function ameliorates the brood size decline seen in *gcna-1* null mutant animals, the catalytic domain and C-terminus of the protein are required to prevent genome instability across generations. Together with the fly experiments, these data indicate that GCNA is a multi-functional protein with the IDR and SprT domain governing distinct aspects of genome integrity.

### Loss of *GCNA* correlates with genomic instability in human germ cell tumors

Our work in flies, worms, and zebrafish indicates that GCNA regulates genome stability across species. To test whether the same held true in humans, we carried out sequence analysis of data obtained from human tumor samples. We performed whole-exome and targeted deep sequencing, SNP array, DNA methylation array, and RNA sequencing on a cohort of 233 patients with pediatric germ cell tumors (GCTs, Table S1), and conducted an integrated analysis of the data analyzed (Xu and Amatruda, in preparation). To investigate tumor suppressor genes that are frequently silenced by copy number (CN) loss (based on SNP array data) and promoter hypermethylation (base on DNA methylation array data) in GCTs, we examined 94 of 233 GCTs in our cohort that were processed by both array technologies. In these 94 GCTs, we calculated the percentage with either CN loss, promoter hypermethylation, or both alterations, for *GCNA* and for 441 known cancer genes (from Catalogue of Somatic Mutations in Cancer (COSMIC) database). Interestingly, we found *GCNA* has the highest alteration frequency (66%, 62 out of 94 cases) among all 442 genes studied here (Figure 6A,B), suggesting *GCNA* to be a top candidate for a tumor suppressor involved in GCT development. No somatic protein-altering mutation was found in *GCNA* gene through our whole-exome and targeted deep sequencing analysis in 137 GCT cases.

**Figure 6.**
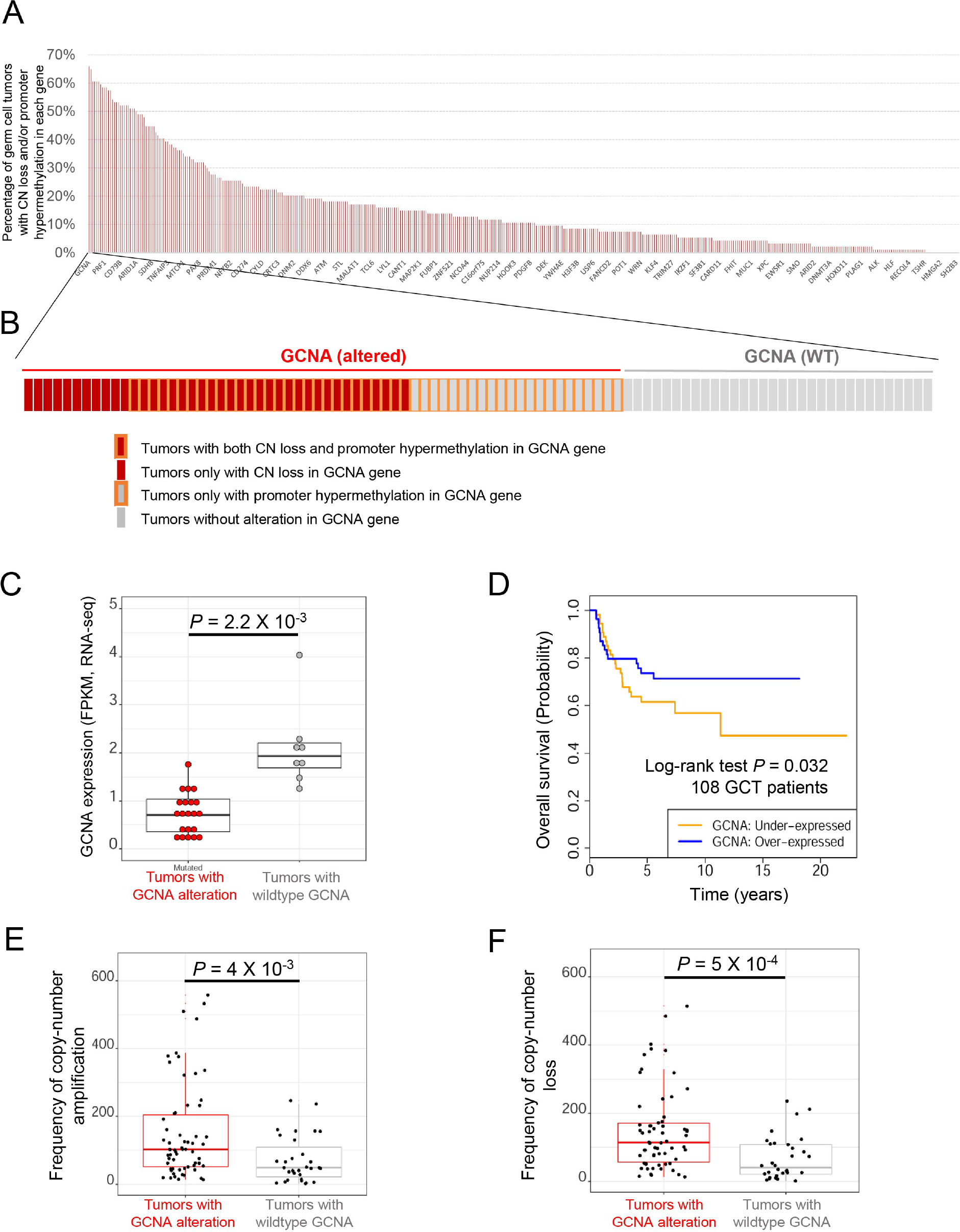
*GCNA* is frequently altered in pediatric germ cell tumors (GCTs), and is associated with poor patient survival and genomic instability. **(A)** *GCNA* genes display the most frequent copy-number loss and promoter hypermethylation in 94 childhood patients with GCTs. **(B)** Frequency of copy-number loss and promoter hypermethylation of *GCNA* gene in pediatric GCTs. **(C)** Copy-number loss and promoter hypermethylation of *GCNA* is associated with low GCNA expression in pediatric GCTs. **(D)** Low GCNA expression is statistically significantly associated with poor survival of GCT patients. **(E)** Copy-number loss and promoter hypermethylation of GCNA associates with copy number amplification across the genome. **(F)** Copy-number loss and promoter hypermethylation of GCNA associates with copy number loss across the genome.

As expected, CN loss and/or promoter hypermethylation in *GCNA* was correlated with significantly lower *GCNA* expression in tumors, compared to GCTs without alterations (Figure 6C). Thus, it appears that genetic and epigenetic alterations are responsible for down-regulation of *GCNA* expression in human GCTs. Notably, we observed a significant association between low *GCNA* expression and poor GCT patient survival (Figure 6D, hazard ratio = 0.66, 95% CI 0.45-0.96, log-rank test, *P* = 0.032), which suggests that *GCNA* might serve as a potential prognostic marker.

To further investigate whether genetic and epigenetic alterations in *GCNA* gene are associated with genome instability, we studied the frequency of copy number amplification and loss on a genome-wide scale. We found that tumors with alterations in *GCNA* gene display significantly elevated frequency of both copy number amplification and loss events (Figure 6E), supporting the idea that *GCNA* expression may contribute to human germ cell tumorigenesis by regulating genome stability.

## DISCUSSION

Here, we report the initial functional characterization of GCNA as a key regulator of genome stability within germ cells and early embryos. *GCNA* mutants exhibit remarkably similar phenotypes in flies, worms, zebrafish, and humans indicating that GCNA carries out conserved functions throughout the animal kingdom. Germ cells must contend with many distinct challenges imposed by their unique biology and the dangers of various endogenous and exogenous genotoxic threats. Loss of GCNA compromises the ability of germ cells to handle these stresses and disrupts a number of seemingly disparate processes including germ cell development and maintenance, chromosome segregation, the cell cycle, DNA replication, and the formation of programmed double stranded breaks during meiosis. Our findings suggest that these processes may be linked together through *GCNA* in unexpected ways. Given its enriched germline expression and critical function, GCNA should be considered, alongside with Nanos, Vasa, and Piwi as an essential player in germ cell biology.

How can GCNA influence so many different processes within germ cells? Our data indicate that the phenotypes exhibited by *GCNA* mutants do not extend from one specific malfunction, but rather from disruption of a number of distinct functions. GCNA contains an N-terminal acidic IDR, followed by a SprT domain, ZnF and HMG box. Our genetic experiments in flies and worms indicates specific functions can be attributed to distinct domains (Figure 5). For example, SprT enzymatic activity appears necessary for the prevention of excessive Spo11-dependent and -independent DNA damage during *Drosophila* oogenesis. By contrast, SprT mutant transgenes rescue the maternal-effect semi-lethality of *GCNA* mutants, indicating this enzymatic activity is dispensable for the proper regulation of chromosome segregation and cell cycle during early embryogenesis. Similarly, in *C. elegans*, *gcna-1* alleles which encode just the amino-terminal IDR domain do not exhibit a reduction in average brood size over the first eight generations. This contrasts with the loss of fecundity conferred by the null allele, again indicating that many germ cell activities depend on the IDR of GCNA. This separation of structure and function provides an important framework for further understanding how GCNA ensures genomic stability across species. Of note, the mouse protein is comprised of just an IDR domain, raising the possibility that the chromosome segregation and cell cycle functions may represent the core activities of these proteins.

Given the modest increase of DPCs within *GCNA* mutants (Figure 4) and the failure of the HE>AA mutant transgene to rescue Spo11-dependent and -independent damage in *Drosophila* ovaries (Figure 5), it is tempting to speculate that the SprT domain of GCNA serves the same function as it does in Spartan proteins, namely to regulate DPCs that form on chromosomes. While GCNA does not contain a PIP (PCNA-Interacting Protein) domain, which mediates interactions between Spartan and PCNA, it nevertheless appears to associate with replication machinery, based on its ability to immunoprecipitate components of the MCM complex (Figure 4). *GCNA* mutants likely suffer from replicative stress based on increased RPA foci in *Drosophila* cells (Figure 3), microsatellite instability in worms (Figure 3), and copy number amplification and loss in human germ cell tumors (Figure 6). This raises the question regarding the separate requirements for GCNA and Spartan. Germ cells may have an increased DPC load compared to somatic cells, and therefore require alternative mechanisms for dealing with these lesions. For example, germ cells undergo extensive epigenetic reprogramming. These reactions, such as histone demethylation, produce cross-linking agents as by-products (Stingele et al., 2017). In addition, germ cells encounter enzymatic DPCs during meiotic DSB induction when Spo11 acts through a topoisomerase-like mechanism and becomes covalently attached to DNA at the break site. While the MRN complex clears these adducts through endonuclease cleavage, perhaps GCNA represents an alternative mechanism for clearing Spo11 off of meiotic chromosomes in order to ensure that all lesions are repaired prior to embryogenesis. Given the increase of Spo11-induced breaks in flies, where very few meiotic breaks are made, GCNA may have evolved to limit either Spo11 activity or the amount of Spo11 that reaches the DNA, perhaps through sequestration of Spo11 or its accessory factors in the cytoplasm. The dramatic accumulation of Topoisomerase 2 in a subset of nuclei in *gcna-1* mutant worms and on mitotic DNA in mutant flies is also consistent with a sequestration model. Lastly, germ cells, particularly oocytes, undergo cell cycle for long periods of time. These cells may accumulate DPCs that would otherwise interfere with chromatin-based processes upon fertilization. Perhaps GCNA provides a replication-independent means for clearing or preventing such lesions.

IDR proteins have emerged as important regulators of germ cell biology. Many proteins that play essential roles in germ cells, such as Vasa, Oskar, Bucky ball and MEG-3, contain IDRs. These IDRs often control the ability of these proteins to undergo phase transitions, allowing for the compartmentalization of various RNAs and proteins. We are just beginning to understand the functional significance of this molecular behavior. The IDR of GCNA is essential for its function within germ cells. While the primary amino acid sequence of GCNA’s IDR has diverged, this region has continued to retain a high percentage of aspartic acid and glutamic acid residues. The functional significance of this distinct composition remains unknown, but perhaps it regulates the stability of GCNA or its ability to form condensates. Our protein expression analysis indicates that GCNA localizes in a discrete particulate pattern in the cytoplasm and on chromosomes during mitosis. In flies and worms, the IDR of GCNA is essential for proper chromosome segregation and cell cycle regulation. Perhaps GCNA undergoes phase transitions and thereby controls the availability, assembly, or function of factors needed for these processes. Indeed, we observe a redistribution and accumulation of Top2 on chromosomes in the absence of *GCNA*.

In flies and zebrafish, loss of *GCNA* leads to profound maternal-effect phenotypes, marked by chromosome segregation defects, chromosome bridges, and cell cycle asynchrony. Worm *gcna-1* mutants also exhibit an increase in chromosome bridges and X-chromosome loss, albeit at a lower frequency or only in later generations. The chromosome bridges that form in embryos derived from *Drosophila GCNA*^*KO*^ females, often contain sequences corresponding to the heterochromatic 359-Bp repeats found on the X chromosome. Interestingly, the 359-Bp satellite is responsible for the chromatid separation defects that occur in hybrid progeny between *D. melanogaster* and *D. simulans* (Ferree and Barbash, 2009). In addition, recent work has shown that loss of *mh*, which encodes the *Drosophila* homolog of Spartan, also results in chromosome bridges that contain 359-bp sequences (Tang et al., 2017). While, our genetic experiments indicate that GCNA and Spartan proteins act in parallel pathways, they may both converge on a specific mechanism that helps to resolve segregation defects involving heterochromatic sequences. Interestingly, we observe the formation of micronuclei and chromosome fragmentation in both *Drosophila* and *C. elegans GCNA* mutants. Similar events are thought to presage chromothripsis in cancer cells. Sequencing data in the accompanying paper indicate that *C. elegans GCNA* mutants display molecular signatures consistant with chromothripsis. Thus, the further study of GCNA may provide a model for understanding the origins of this newly recognized process and for determining how germ cells protect themselves from widespread chromosome re-arrangements in the face of DNA damage.

Loss of GCNA results in much more severe phenotypes in flies and zebrafish, relative to worms. We still do not fully understand the basis of these differences. Perhaps, worms have evolved redundant mechanisms that compensate for the loss of GCNA activity or their unique holocentric chromosome structure avoids potential problems experienced by organisms that rely on centromeres for proper chromosome segregation. Interestingly, *Drosophila* encodes for three GCNA orthologs (Figure 1; S1), all of which exhibit enriched expression in gonads based on modEncode data (http://flybase.org). GCNA (CG14814) exhibits enriched expression in ovaries, GCNA2 (CG2694) exhibits enriched expression in the ovary and testis and GCNA3 (CG11322) exhibits enriched expression in the testis. The expression of GCNA2 significantly increases in the absence of GCNA (RNA-seq data; Figure S3), suggesting that feedback regulatory loops may control the expression of these genes. Further experiments will be needed to test for potential functional redundancy.

Germ cells have many unique features in regards to reprogramming, regulation of the cell cycle and DNA repair. The study of *GCNA* will provide further insights into how these processes are coordinated with each other to ensure the faithful transmission of genetic material from one generation to the next. In addition to the observed correlation between loss of *GCNA* and human germ cell tumors, we anticipate *GCNA* function likely influences other aspects of human fertility and transgenerational inheritance across sexually reproducing species.

## ACKNOWLEDGEMENTS

We would like to thank K. McKim, T. Orr-Weaver, D. Ardnt-Jovin, M. Osterfield, K. Dubrovinski, M. Lilly, J. Abrams, H. Richardson, the Bloomington *Drosophila* Stock Center and the Developmental Studies Hybridoma Bank for *Drosophila* reagents and Sarit Smolikove and Aimee Jaramillo-Lambert for worm antibodies. Some strains were provided by the CGC, which is funded by NIH Office of Research Infrastructure Programs (P40 OD010440). We also thank Diwash Jangam and Esther Bertran for their help with analysis of repeat elements in RNA Sequencing data. We thank members of the Buszczak and Yanowitz labs for comments and advice, and Jose Cabrera for help in preparing the graphical abstract. V.B. was supported by HHMI (Grant#56006776), NIGMS (T32GM10977601) and a fellowship from the Center for Regenerative Science and Medicine at UT Southwestern. C.D.G. was supported by NIGMS (5T32GM008203-30). This work was also supported in various phases by awards from the NIGMS (R01GM086647) and the NIA (R01AG047318) to M.B. and by NIGMS (R01GM104007) to J.L.Y. J.F.A. was supported by grant RP110394 from the Cancer Prevention and Research Institute of Texas and a Consortium Research Grant from the St. Baldrick’s Foundation.

## MATERIALS and METHODS

### Fly stocks

Fly stocks were maintained at 20°C–25°C on standard cornmeal-agar-yeast food. nanos-gal4-VP16 was a gift from Y. Yamashita. *mei-w68*^*0949*^ and *mei-P22^1^* stocks were a gift from K. McKim. p53-GFP reporter was a gift from J. Abrams. *mh*^*1*^ (BL# 7630), *Histone2Av.mRFP* (BL# 23651) and *Df(1)BSC719* (GCNA Df; BL# 26571) stocks were obtained from Bloomington Stock Center, Indiana.

### Immunofluorescence

Adult ovaries were stained according to Tastan et al. (2010). *Drosophila* embryos were stained according to S.R. Mani et al. (2014). The following primary antibodies were used: mouse anti gamma-H2AV (DSHB unc93.5.3.1) (30:200), rabbit anti-gamma-H2Av (K. McKim) (1:500), rabbit anti-C(3)G (Mary Lilly) (1:3000) (Hong et al., 2003), rabbit anti-RPA (1:500) (Terry Orr-Weaver from Fisher and Cotterill labs), rabbit anti-Top2 (T. Hsieh and D. Ardnt-Jovin) (1:400), mouse anti-HTS (1B1) (DSHB) (1:20), rat anti-Vasa (DSHB) (1:20), rabbit anti-GFP (1:1000) (Life Technologies), rat anti-HA 3F10(Roche), and fluorescence-conjugated secondary antibodies (Jackson Laboratories)(1:300).

### Western Blot analysis

Proteins extracts were separated by SDS-PAGE and transferred onto a nitrocellulose membrane. The following primary antibodies were used: rabbit anti-Top2 (T. Hsieh and D. Ardnt-Jovin) (1:2000), rat anti-Vasa (DSHB)(1:200), mouse anti-actin (DSHB) (1:100), rat anti-HA 3F10 (Roche)(1:2000). The secondary antibodies were anti-mouse IgG HRP (Jackson ImmunoResearch) (1:2000), anti-rabbit IgG HRP(Jackson ImmunoResearch) (1:2000), and anti-rat IgG HRP(Jackson ImmunoResearch) (1:2000).

### DNA-protein Crosslink Isolation

DPCs were isolated and detected using a modified rapid approach to DNA adduct recovery (RADAR) assay wherein the tissue was lysed under denaturing conditions and the DNA was ethanol precipitated (Kiianitsa and Maizels, 2013; Vaz et al., 2016). For *Drosophila* the ovaries were dissected and lysed for RADAR and the embryos were 0-2hrs old when lysed. The zebrafish embryos were lysed at the 1000 cell stage.

### DNA and DNA-protein Crosslink Detection

DNA was detected using a slot blot vacuum manifold (Biorad). Specific proteins were detected by normalizing to DNA concentration and digesting with benzonase. The proteins were separated on a polyacrylamide gel. The following primary antibodies were used: mouse anti-dsDNA (abcam) (1:2000) and rabbit anti-Top2 (T. Hsieh and D. Ardnt-Jovin) (1:2000).

### Generating the *GCNA*^*KO*^ alleles

To generate the *GCNA*^*KO*^ allele, guide RNAs were designed using http://tools.flycrispr.molbio.wisc.edu/targetFinder and synthesized as 5-unphosphorylated oligonucleotides (Table S3), annealed, phosphorylated, and ligated into the BbsI sites of the pU6-BbsI-chiRNA plasmid (Gratz et al., 2013). Homology arms were PCR amplified and cloned into pHD-dsRed-attP (Gratz et al.,2014) (Addgene). Guide RNAs and the donor vector were co-injected into nosP Cas9 attP embryos at the following concentrations: 250 ng/ml pHD-DsRed-attP donor vector and 20 ng/ml of each of the pU6-BbsI-chiRNA plasmids containing the guide RNAs (Rainbow Transgenics).

### Cloning of *Drosophila GCNA* transgene

PCR products were cloned into pENTR (Life Technologies) and swapped into pAHW, pAWG (attB added by Tony Harris) or pAFHW (Drosophila Gateway Vector Collection) using an LR reaction. Using this approach we isolated clones corresponding to the CG14814 cDNA (accession number BDGP:RE06257) transcripts.

### Live Imaging

3-5 day old males and virgin females were mated in mating cages containing grape juice (3%) agar plates with a little bit of wet yeast. The flies were allowed to lay eggs for 1-2 hrs at 25 C. Eggs were carefully collected and dechorionated by rolling them on double-sided tape pasted on a slide. Dechorionated eggs were then mounted using oil and non-auto-fluorescent glue. Live imaging was conducted every 15 seconds using a Zeiss LSM800 microscope.

### Fertility Assays

0-2 day-old males and virgin females of the appropriate genotype were mated in mating cages with grape juice (3%) agar plates with a little bit of wet yeast. The flies were allowed to lay eggs for 48-72 hrs at 25°C before switching out the plates. Flies were allowed to lay eggs for a total of 7 days before eggs were counted.

### Hydroxyurea exposure

0-2 day-old wildtype and *GCNA*^*KO*^ female flies were collected and fed on wet yeast for 24 hrs. They were then starved for 16-18 hrs. Whatman paper was soaked in either water +2% sucrose or 50mM Hydroxyurea. Flies were allowed to feed on sucrose with solvent or drug for 24 hours before being dissected and immunostained.

### Super-Resolution Imaging Structured illumination microscopy (SIM) imaging and image processing

Ovaries were dissected according to standard protocol and mounted in Prolong Gold antifade reagent (Life technologies). Nikon N-SIM system (Nikon, Tokyo, Japan) equipped with a 100×/1.49 TIRF oil immersion objective lens (Nikon), the iXon + electron multiplying charged-coupled device camera (Andor) and an excitation laser unit of 405 nm, 488 nm, 561 nm and 640 nm (Coherent) was used for a super-resolution optical imaging. Z-stacks of SIM optical sections were acquired with a 120 nm Z-step size. Image processing, including 3-dimensional reconstruction and co-localization analysis, were carried out using the NIS-Element Advanced Research software (Nikon).

### Electron Microscopy

Eggshells were mounted on SEM stubs and sputter coated with gold/palladium in a Cressington 108 auto sputter coater. Images were acquired on a Field-Emission Scanning Electron Microscope (Zeiss Sigma, Carl Zeiss, Thornwood, NY) at 10.0 kV accelerating voltage.

## FISH

A protocol adapted from the Fox lab was used with oligopaints (Beliveau et al., 2014; Beliveau et al., 2015; Beliveau et al., 2012). Oligos with fluorophores were ordered from Integrated DNA technologies.

### RNA Sequencing

30 ovaries were dissected from GCNA^KO^ and control (*w*^*Berlin*^) flies. There were three biological replicates produced from independent backcrosses of the KO into *w*^*Berlin*^ background for each sample. Qiagen miRNeasy Mini kit was used for RNA extraction and the RNA was sent for library preparation that selected for total RNA at McDermott Sequencing core at UTSW. The samples were sequenced in Illumina HiSeq 2500 cycle single-read platform. The sequencing reads were checked for quality using FastQC program (http://www.bioinformatics.babraham.ac.uk/projects/fastqc/). For analyzing differential gene expression, STAR aligner (version 2.5) (Dobin et al. 2013) was used to map the RNA-Seq reads to the *Drosophila* reference genome (genome assembly BDGP6.88) with an additional flag --outFilterMultimapNmax 100. TEtranscripts (Jin et al. 2015) was used to generate the read counts that mapped to TEs and genes. DESeq2 (Love et al. 2014) was used for differential expression analyses of TEs with a FDR of 5%.

### *C. elegans* stock maintenance

Worm strains were established and maintained at 20°C for long-term passaging under standard conditions (Brenner, 1974). For all experiments, unless otherwise stated, F1 homozygous *gcna-1(ea43)* worms were shifted as L4/young adults to 25°C and grown for two generations before analysis, since brood analysis showed that F1 and F2 populations sizes were near wild-type, suggesting maternal rescue. Wild type refers to the *C. elegans* variety Bristol, strain N2. The stocks utilized in this study were: *prg-1(n4357)* I, *dog-1(gk10)* I, *xpf-1(e1487)* II, *gcna-1(ea43*) III/ <u>hT2</u> [<u>bli-4</u>(<u>e937</u>) <u>let-?</u>(<u>q782</u>) <u>qIs48</u>] (I;III), *gcna-1(ea67)*[*ollas::gcna-1*]) III, *gcna-1(*ea72 [*HA*::*gcna-1*]) III, *gcna-1(ea76)* III/ <u>hT2</u> [<u>bli-4</u>(<u>e937</u>) <u>let-?</u>(<u>q782</u>) <u>qIs48</u>] (I;III), *gcna-1(ea80) exo-1(tm1842)* III/ <u>hT2</u> [<u>bli-4</u>(<u>e937</u>) <u>let-?</u>(<u>q782</u>) <u>qIs48</u>] (I;III), *rfs-1(ok1372)* III/ <u>hT2</u> [<u>bli-4</u>(<u>e937</u>) <u>let-?</u>(<u>q782</u>) <u>qIs48</u>] (I;III), *gcna-1(ea81*) *rfs-1(ok1372)* III/ <u>hT2</u> [<u>bli-4</u>(<u>e937</u>) <u>let-?</u>(<u>q782</u>) <u>qIs48</u>] (I;III), *spo-11(ok-79)* IV*/ nT1 [unc-?(<u>n754</u>) let-? qIs50] (IV;V)*, *dvc-1(ok260)* V, *dvc-1(ea65)/nT1*[qIs51] V, *sws-1(ea12)* V, *prg-1(n4357)* I;*gcna-1(ea43)* III, *xpf-1(e1487)* II; *gcna-1(ea43)* III, *gcna-1(ea43)* III;*spo-11(ok-79)* IV*/ nT1 [unc-?(<u>n754</u>) let-? qIs50]* (IV;V), *gcna-1(ea43)* III; *dvc-1(ea65)* V, *gcna-1(ea43)* III;*sws-1(ea12)* V. Primers and PCR conditions for genotyping are provided in Table S2. Several of the stocks were provided by the *Caenorhabditis* Genome Center (University of Minnesota) which is funded by NIH Office of Research Infrastructure Programs (P40 OD010440).

### Generation of *C. elegans* CRISPR alleles

Null, tagged, and mutated versions of *gcna-1* were created by CRISPR/Cas9 genome engineering. For null alleles *ea43* and linked double mutant lines, 5’ *gcna-1* crRNA TCGAAATGGTGTAGGCATTG and 3’ *gcna-1* crRNA CTAAAACATTCGGATGTGAT were utilized with repair ultramer: 5’ ttctaacttttaagttgatattcagaatATCGTGCTCACTGATCAGCAACGACCAGATACAACGATTGATCAATCGGAAGATCCTGTAGAGGAAAAAGAGGATGAT 3’. For *ea76*, internal crRNA AACTGTGTCATGCAGCAACA was used with the 5’ *gcna-1* crRNA. For tagging with OLLAS, the ultramer sequence was: TTTTCTCGTATTCTGCAAATCTTCTGTATTCCTCAATGTCCGGATTCGCCAACGAGC TCGGACCACGTCTCATGGGAAAGGGAGGAGGAGGAGGACCTACACCATTTCGAGATCTTCATAACAAAAGTAA. For HA tagging, the ultramer used was: TTTTCTCGTATTCTGCAAATCTTCTGTATTCCTCAATGTACCCATACGATGTTCCAGATTACGCTGGAGGAGGAGGAGGACCTACACCATTTCGAGATCTTCATAACAAAAGTAA. The *dvc-1(ea65)* null was created with 5’ *dvc-1* crRNA GAGCTGCGACTCATtctttg and 3’ *dvc-1* crRNA GACATCTGGATTACTGTCTC and repair template ultramer: CTA CTC GAT TTC CAC ATC TCC ATC GTT CAC ATA ATT TCC GCA GCC ACA AAT CTC GGG TGA AGC TAT CTA TAC TCA TGT AAT TTT TAT TAT GTA CTA CAC T.

Injection mixes containing crRNA, tracrRNA, ssDNA ultramers, and Cas9 protein were prepared as described (Paix et al., 2017). crRNA and ssDNA repair template for *dpy-10(cn64)* co-injection marker were also included (Arribere et al., 2014). Both Roller and Dumpy (*dpy-10*) transformants were individually plated and F2 Dumpy progeny were screened by PCR for relevant insertions and deletions (see Table S2 for primers and conditions). Putative mutants were outcrossed to N2 at least twice prior to sequencing.

### Fertility Assays

Brood sizes were performed by individually plating L4 animals on center-seeded 3cm plates and transferring every 24 hours until the cessasion of egg-laying. Total numbers of L4 and adult hermaphrodite and male offspring were counted 48-72 hours later.

Transgenerational assays were performed by starting 3cm or 12-well plates with single F1 progeny of balanced heterozygous moms. Each generation, the first F1 progeny to reach L4 were transferred. Plates were examined after 24 hours to determine if the worm was sterile and replaced with an adult from the parental plate if no eggs were observed in the uterus or on the plate. Brood sizes were binned into ranges shown in Figure 1. Presence of males and mutant animals (Dumpy, Uncoordinated, etc) was noted. Sterile populations were reassessed by plating additional worms from the parental plate and classified as sterile when no further offspring could be attained from the previous generation. This methods ensures that sterility is inherent to the population and not simply a subset of offspring. All experiments utilized non-starved populations of worms. Data is shown for 25°C as sterility did not arise in populations of *gcna-1(ea43)* grown at 20°C.

Fecundity of *gcna-1;prg-1* hermaphrodites was tested by shifting *gcna-1(ea43)*, *prg-1*, or *prg-1;gcna-1(ea43)* animals from 20°C to 25°C as early L4 larvae and counting offspring 3 - 4 days later.

### Detecting microsatellite deletions

The *dog-1(gk10)* mutations was outcrossed to N2 twice before mating with *gcna-1(ea43)*. *dog-1, dog-1;gcna-1*, and *gcna-1* lines were generated from the cross and grown three generations at 25°C. For each genotype 100-200 F4 worms were collected in 14 μl of lysis buffer (50 mm KCL, 10 mm, Tris pH 8.3, 2.5 mm MgCl_2_, 0.45% NP-40, 0.45% Tween-20, 0.01% gelatin) with 5 mg/ml freshly added proteinase K and and individually lysed at 60° for 60 min and then at 95° for 15 min. Deletions upstream of the *vab-1* locus were detected using a nested PCR reaction as described (Youds et al., 2006). PCR products were run on a 2-3% agarose gel and scored for unique DNA fragments less than 500bp.

### Irradiation sensitivity assay

Day 1 adults were exposed to increasing doises of ionizing radiation using a ^137^Cs source (Gammacell 1000 Elite; Nordion International). Embryos and L1 larvae from individually-plated animals at t= 24-36 hours post-irradiation were collected and counted. Viable offspring were counted 2.5 – 3 days later. Data is normalized to the hatching rates of the unexposed animals of each genotype.

### Quantitative PCR

Approximately 100 young adults of a each genotype were washed three times in 1x M9 buffer (3 g/L KH_2_PO_4_, 6 g/L Na_2_HPO_4_, 5 g/L NaCl, 1 mM MgSO_4_), resuspended in Trizol (Invitrogen), and vortexed for ~60 seconds before being flash frozen and stored at − 80°C. Once all the samples were collected, samples were thawed on ice, sonicated, and RNA was isolated by chloroform extraction and isopropanol precipitation. Samples were treated with DNase (Sigma #AMPD1) and reverse transcribed into cDNA (Protoscript m-MuLV First Strand cDNA Synthesis kit, NEB #E6300S) according to manufacturer’s instructions. Quantitative real-time PCRs were performed on the Applied Bio Systems 7300 Real Time PCR System using Sybr Green chemistry (SensiMix SYBR Hi-ROX kit, Bioline #QT-605) with transcript-specific primers for Tc1, Tc2, Tc3, and Tc4v as described in Table S2. The reference genes *rpl-32* (Hoogewijs et al., 2008) was used for normalization across samples and gene expression was analyzed using the ΔC_T_ method (Livak and Schmittgen, 2001). Results are presented as the average of combined data from three independent biological replicates that in turn is comprised of three technical replicates each.

### *C. elegans* immunostaining

One day-old adult worms were dissected in 3.5µL 1x sperm salts (50mM PIPES, pH 7.0, 25 mM KCL, 1 mM MgSO_4_, 45 mM NaCL, 2 mM CaCl_2_.) + 0.2µL 10mg/mL levamisole. Fixation and pre-hybridization varied for different primary antibodies:

α-RAD-51. 7µL of 2% paraformaldehyde was added to slides and incubated 5 minutes in a humidity chamber at room temperature before freezing on a metal block on dry ice for 10 minutes. Coverslips were flicked off and slides were then submerged in −20°C methanol for 5 minutes and dipped in −20°C acetone for 5 seconds. Slides were air dried, a wax box was drawn around the samples, and dissected worms were prehybridized for 3x 10 minutes in Phosphate buffered saline (PBS) + 0.1% Tween 20 +0.1% BSA (PBSTB). Primary antibody (courtesy of Sarit Smolikove) was diluted 1:20,000 and incubated overnight at 4°C.

α-FLAG (for visualization for TOP-2::FLAG). 3.5µL of 2% Triton and 7µL 2% PFA were added to dissected samples (fixed and frozen as above). Slides were submerged in − 20°C methanol for 1 minute and washed as above in PBSTB. Mouse α-FlagM2 (Sigma F1804) was diluted 1:500 and incubated overnight at 4°C.

α-HA (for visualization of HA::GCNA-1) 3.5µL of 2% Triton and 7µL 2% PFA were added to dissected samples (fixed and frozen as above). After fixation, slides were submerged in −20°C Methanol for 5 minutes. Mouse α-HA antibody (Santa Cruz F7) was diluted 1:500 and incubated overnight at 4°C.

The next day, slides were washed 3 x10 min each with PBSTB and then incubated with secondary antibodies diluted in PBSTB (goat α-Rabbit Alexa 568, goat anti-mouse Alexa 488, diluted 1:2000) for 2 – 4 hours at room temperature. Slides were then washed once with PBSTB for 10 min, once with PBS+DAPI (4’,6-diamidino-2-phenylindole) for 15 min, and again with PBSTB for at least 10 min prior to mounting in Prolong Gold Antifade Mountant with DAPI (ThermoFisher Scientific, P36931). Slides were cured overnight prior to confocal imaging.

### Whole-mount staining for diakinesis analysis

One-day old adult worms were fixed in Carnoy’s solution (three parts absolute ethanol; two parts chloroform; one part glacial acetic acid), stained with DAPI (4’,6-diamidino-2-phenylindole) for at least fifteen minutes, then mounted in Prolong Gold Antifade Mountant with DAPI (ThermoFisher Scientific, P36931). Slides were cured overnight prior to confocal imaging. Statistical analyses were performed as described in Macaisne et al., 2018 using GraphPad Prism software.

### Imaging and Quantification of Staining

RAD-51 foci and diakinesis nuclei were quantified by collecting Z-stack images on a Nikon A1r confocal microscope with 0.2µm sections. Three dimensional stacks were visualized using Volocity imaging software (Quorum Technologies). For RAD-51 foci, nuclei from the transition zone nuclei through the pachytene/ diplotene border were quantified. The pachytene region was divided into 6 equal parts according to number of rows of nuclei in this region. Diakinesis nuclei were rotated in three dimensions to attain the number of DAPI-staining bodies in the −1 and −2 nuclei (i.e. the two oocytes preceding the spermatheca).

### Zebrafish CRISPR

Target sites for sgRNAs were selected using the online software CRISPR DESIGN (http://crispr.mit.edu). One sgRNA was designed to target a site on exon 3 (5’-GAAGACCAGACGTCCAGCTT-3’) of the zebrafish *gcna* gene. The sgRNA was synthesized as previously described (Gagnon et al., 2014). Briefly, the gene-specific oligonucleotide, consisting of an upstream SP6 promoter (5’-GCGATTTAGGTGACACTATA-3’) followed by the 20-base target sequence (5’-GAAGACCAGACGTCCAGCTT-3’) and a sequence complementary to the reverse tracrRNA tail oligonucleotide (5’-GTTTTAGAGCTAGAA-3’), was annealed to the reverse tracrRNA tail oligonucleotide, followed by incubation with T4 DNA polymerase to fill the ssDNA overhangs. The resulting DNA template was then purified using QIAquick PCR Purification Kit (Qiagen) and used for sgRNA transcription using the Megascript T7 Kit (Ambion). The sgRNA was then treated with DNase and precipitated with LiCl/ethanol.

### Microinjections

Zebrafish embryos were injected at the one-cell stage with 2nl of a mix consisting of sgRNA (83 ng/ul), 1.2ul Cas9 protein (500 ng/ul) (PNA Bio Inc) and 0.08% phenol red dye. The embryos were injected with the PLI-90A picoinjector (Warner Instruments).

### DNA extraction and PCR genotyping

Genomic DNA was extracted from either a pool of three 24hpf larvae or a single caudal fin of adult zebrafish. Tissue was incubated in 50 ul 50mM NaOH at 95°C for 30min. 10ul Tris-HCl (pH 8.0) was added to the lysate and vortexed to neutralize it. 1ul of the lysate was used for each PCR reaction with the forward (5’-GCTTAGGATCGGTAGTTTTCCG-3’) and reverse (5’-GCAGGAGTCCATGTATGGAC-3’) primers. For the PCR reactions the samples were denatured at 95°C for 3 min followed by 40 cycles consisting of 95°C for 30 sec, 55°C for 30 sec and 72°C for 30 sec and a final step at 72°C for 5 min. To identify founders and determine germline transmission of indels, PCR products from embryo lysates were either directly ran on a 3% agarose gel or the T7E1 Assay was performed as described previously (Kim et al., 2009) and the digest products ran on a 1.5% agarose gel. To identify and sequence specific indels in the F1 generation, adult zebrafish PCR products were either sequenced directly with the primers listed above or were cloned into the pCR4-TOPO vector and sequenced with M13 forward and reverse primers.

### Establishment and propagation of *gcna*^*KO*^ zebrafish lines

Adult mosaic fish were out-crossed to wild-type AB Tg(*piwil1:eGFP*) fish to genotype embryos and identify fish with germline transmission. Confirmed founders were subsequently out-crossed to wild-type AB Tg(*piwil1:eGFP*) fish and the progeny was genotyped at adulthood. Heterozygous *gcna* mutants were sequenced and two heterozygous mutants with different indel mutations in the *gcna* gene were mated to generate heteroallelic mutants, with one copy of both mutated alleles. These heteroallelic mutants were in-crossed and also out-crossed to wild-type AB Tg(*piwil1:eGFP*) fish.

### Immunohistochemistry

Embryos were either dechorionated manually post-fixation (< 24hpf) or with Pronase (≥ 24hpf) (Sigma-Aldrich) prior to overnight fixation in 4% paraformaldehyde/1XPBS (PFA) at 4 °C. For DAPI staining, the embryos were deyolked post-fixation incubated for 5 min in DAPI (300 nM DAPI in 0.1% PBS-T) and subsequently washed six times in 0.1% PBS-T. All embryos were mounted with 3% methyl cellulose in E3 medium (5 mM NaCl, 0.17 mM KCl, 0.33 mM CaCl_2_, 0.33 mM MgSO_4_) and images acquired on a Zeiss LSM 800 Confocal Microscope, respectively.

### Embryonic phenotype imaging and classification

To analyze embryonic phenotype, 27 hpf embryos were dechorionated with Pronase, anesthetized with 0.015% MS-222 and mounted in 3% methyl cellulose in E3 medium. Images were taken at 3.2X magnification on a Leica MZ12.5 stereomicroscope attached to a Nikon E4500 camera. Embryos were classified as abnormal if they displayed significant developmental delay in comparison to wild type embryos, as indicated by the ventral body axis curvature posteriorly, the shortened body axis and the presence of dark necrotic tissue.

### Characterization of human tumors

To detect somatic mutations, exome capture was carried out using SureSelect Human All Exon v4+UTRs (Agilent Technologies), and sequencing was performed with a HiSeq 2000 instrument (Illumina) with 100-bp paired-end reads to a mean coverage of 130× for exomes. Raw reads were mapped to human reference genome (hg19) using BWA(Brangwynne et al., 2009). Matched tumor-normal BAM files were used as input for VarScan software (Koboldt et al., 2012) to identify somatic single-nucleotide variants (SNVs) and small-scale insertion/deletions (INDELs).

### Detection of copy-number variation

Genomic DNA from GCT samples was analyzed by SNP array technologies using the Illumina Omni 2.5M SNP array and Affymetrix OncoScan array, according to the manufacturers’ recommendations. We used Nexus Copy Number Discovery 7.0 software (BioDiscovery, Inc.), which can process raw data from both platforms with the same algorithm and procedure. In this software, the data were corrected for GC content and segmented by using SNP-FASST2 algorithm with default parameters.

### Detection of promoter methylation

Genome-wide methylation analysis was performed using the Infinium HumanMethylation450 BeadChip array (Illumina, San Diego, CA) following Illumina’s standard protocol. Raw intensity (idat) files were converted by using the methylumi package(Triche et al., 2013). Combined with IMA package (Wang et al., 2012), DNA methylation sites with missing values, cross hybridizing probes, located within repeat regions or on sex chromosomes were excluded, resulting in a total of 392,714 probes retained. Methylation data were subsequently converted into β values, ranging from 0 (unmethylated) to 1 (fully methylated), and these values were normalized using a beta-mixture quantile normalization method (BMIQ) (Teschendorff et al., 2013).

To detect gene expression and further conduct analysis on association between gene expression and genetic/epigenetic alterations, RNA of GCT samples was sequenced on Illumina HiSeq2000 according to the manufacturer’s protocol (Illumina). 100-bp paired-end reads were assessed for quality and reads were mapped using CASAVA (Illumina). The generated FASTQ files were aligned by Bowtie2 (Langmead and Salzberg, 2012) and TopHat2 (Kim et al., 2013). Cufflinks (Roberts et al., 2011; Trapnell et al., 2010) was used to assemble and estimate the relative abundances of transcripts at the gene and transcript level.

### Survival association analysis

Survival association analysis between gene expression and GCT patients’ survival was calculated based on 108 GCT cases measured by Affymetrix U133A microarray platform (Korkola et al., 2015; Korkola et al., 2009). Signal intensity CEL files were downloaded from Gene Expression Omnibus (GEO) repository at http://www.ncbi.nlm.nih.gov/geo/, data set GSE3218 and GSE10783. CEL files were then processed by Affymetrix Power Tools (APT) with Robust Multiarray Average (RMA) method. Cox proportional hazards model was used to calculate the statistical significance, as well as hazard ratios and 95% confidence intervals of the associations between the gene expression and survival. Kaplan-Meier curves were generated based on gene expression values dichotomized into over-and under-expressed groups using the within cohort median expression value as a cutoff.

---

**Figure S1.**
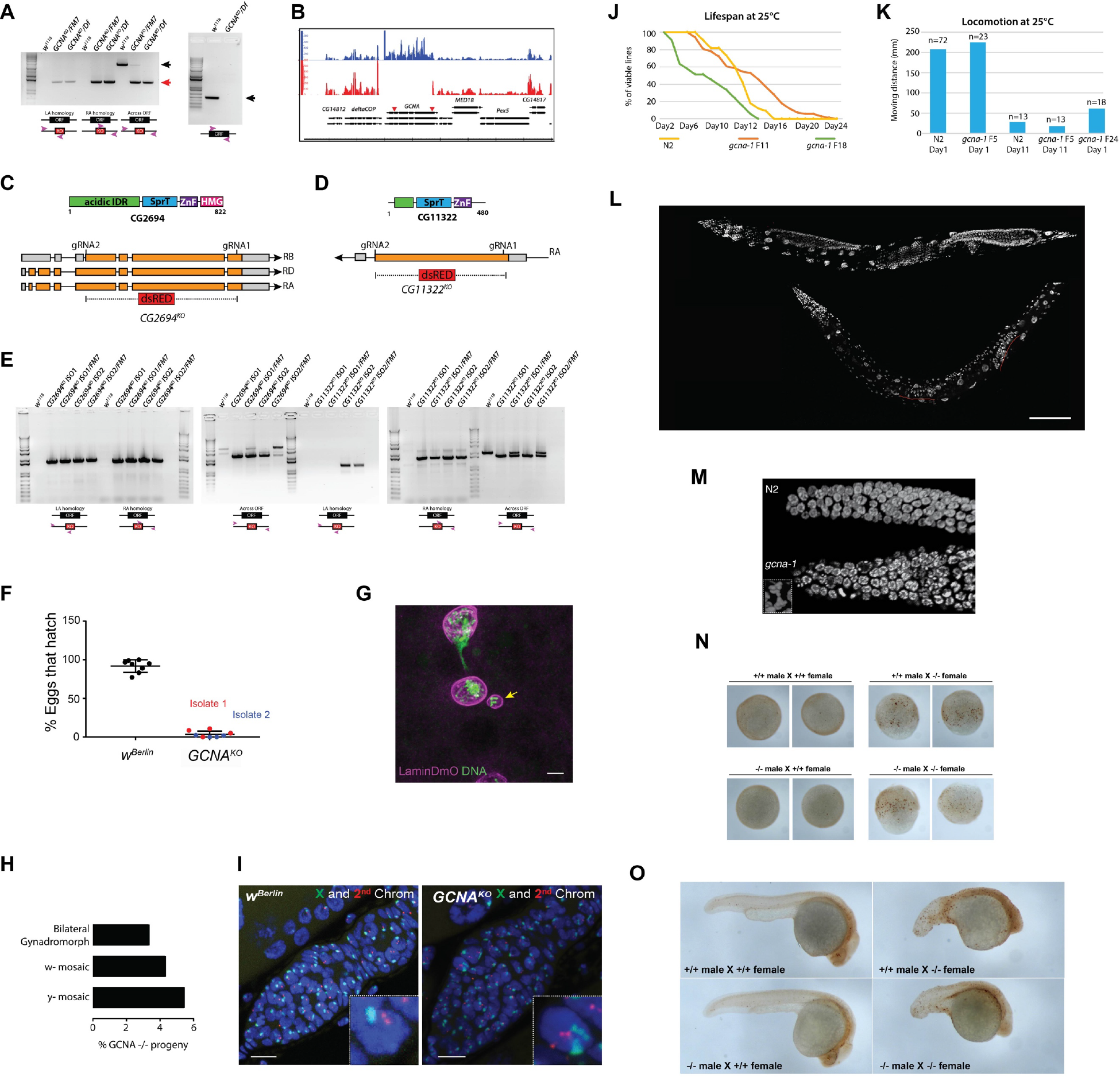
(related to Figure 1) Loss of *GCNA* results in chromosome instability across species. **(A)** PCR verification of the *Drosophila GCNA*^*KO*^ mutation using the indicated primers. **(B)** Reads from RNA-seq showing that *GCNA*^*KO*^ homozygotes do not express GCNA. **(C)** Protein domain structure and gene organization of *CG2694*. **(D)** Protein domain structure and gene organization of *CG11322*. **(E)** PCR analysis of the knockout alleles using the indicated primers. **(F)** Quantification of the percentage of eggs from control and two independent isolates of the *GCNA*^*KO*^ allele that hatch, n>500. **(G)** Image of chromatin bridge and micronucleus in *Drosophila* embryo from *GCNA* mutant mother stained for Lamin Dm (magenta) and DNA (green). **(H)** Quantification of phenotypes observed in progreny of *GCNA*^*KO*^ females that survive to adulthood. **(I)** FISH for the X (green) and second (red) chromosomes performed on control and *GCNA*^*KO*^ mutant ovaries. **(J)** Kaplan-Meier plot of control and *gcna-1* mutant *C. elegans* at indicated generations. **(K)** Quantification of locomotion in control and *gcna-1* mutant *C. elegans*. **(L)** DAPI staining of control and late generation *gcna-1* mutant hermaphrodites showing a reduction in gonad size (red line). **(M)** DAPI-stained germ cells from N2 and late generation *gcna-1* mutant worms showing chromosome bridging in *gcna-1*. **(N)** γH2Av staining of early embryos derived from the indicated crosses. **(O)** γH2Av staining of late stage embryos derived from the indicated crosses.

**Figure S2.**
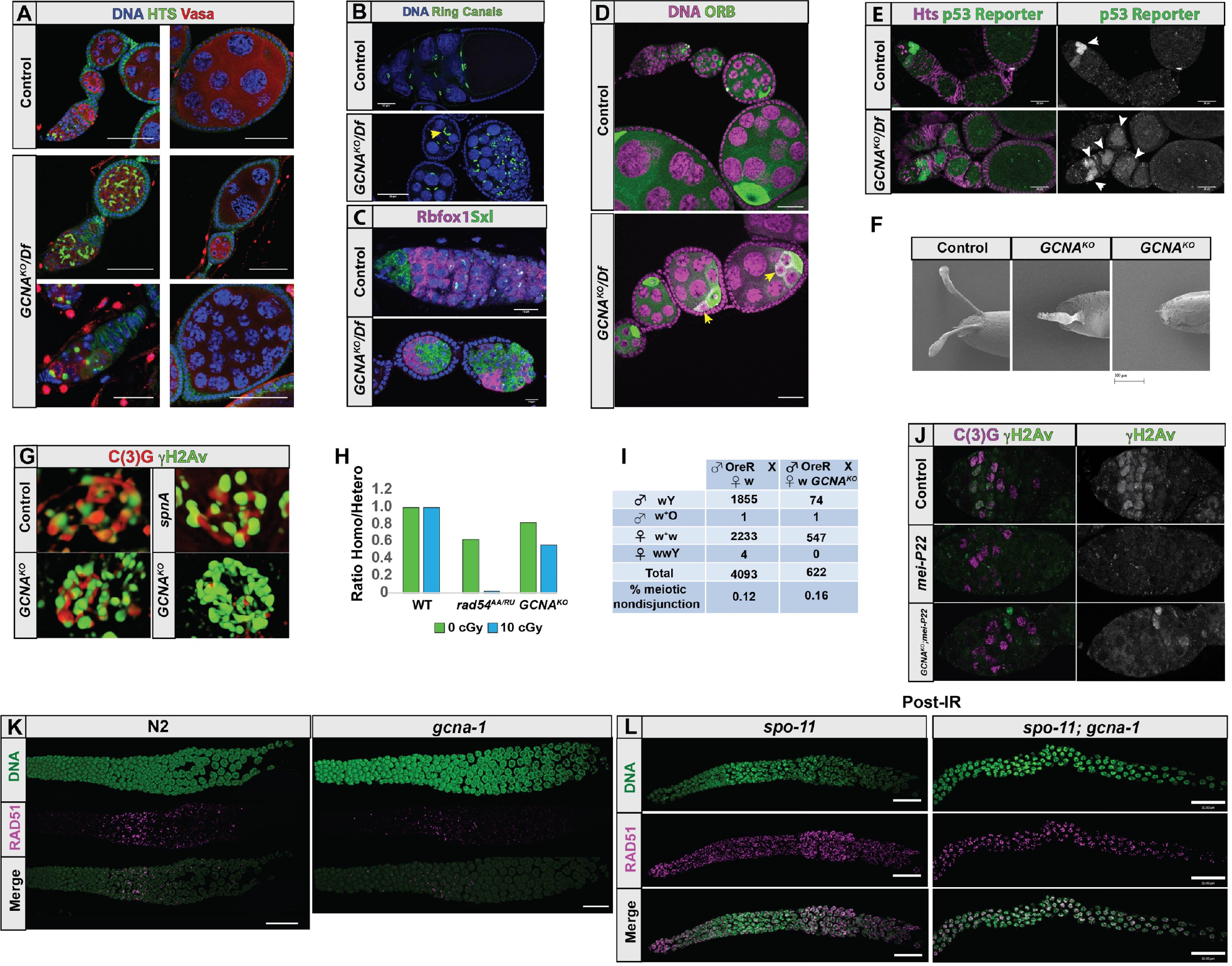
(related to Figure 2). Loss of *GCNA* results in oogenesis defects in *Drosophila*. **(A)** Control and *GCNA*^*KO*^/*DF* ovaries stained for DNA (blue), Hts (green) and Vasa (red). **(B)** Control and *GCNA*^*KO*^/*DF* ovaries stained for DNA (blue) and Hts-RC (green). Yellow arrowhead points to oocyte with only three ring canals. **(C)** Control and *GCNA*^*KO*^/*DF* ovaries stained for Rbfox1 (magenta) and Sxl (green). **(D)** Control and *GCNA*^*KO*^/*DF* ovaries stained for DNA (magenta) and Orb (green). Yellow arrowheads point to germ cells with abnormal accumulation of Orb. **(E)** Control and *GCNA*^*KO*^/*DF* ovaries stained for Hts (magenta) and a p53 reporter (green). **(F)** Scanning Electron Micrograph of control and *GCNA* mutant eggs. **(G)** Structured Illumination Microscopy images comparing control, *spnA*, and *GCNA*^*KO*^ mutant fly meiotic nuclei stained for C(3)G (red) and γH2Av (green). **(H)** Graph showing the ratio of control, *okra (rad54)*, and *GCNA* homozygotes and their heterozygous counterparts that survive to adulthood after receiving a dose of IR. **(I)** Quanitification of genotypes of adult flies that result from crossing control *white (w)* and *w GCNA*^*KO*^ females with OregonR males. *GCNA* mutant females exhibit similar levels of meiotic nondisjunction compared to controls. **(J)** Control, *mei-P22* and *GCNA*^*KO*^; *mei-P22* mutants stained for C(3)G (magenta) and γH2Av (green) **(K)** Control and *gcna-1* mutant worm germ lines stained for DNA (DAPI, green) and RAD-51 (magenta). **(L)** Irradiated *spo-11* and *gcna-1;spo-11* mutant worm germ lines stained for DNA (DAPI, green) and RAD-51 (magenta).

**Figure S3.**
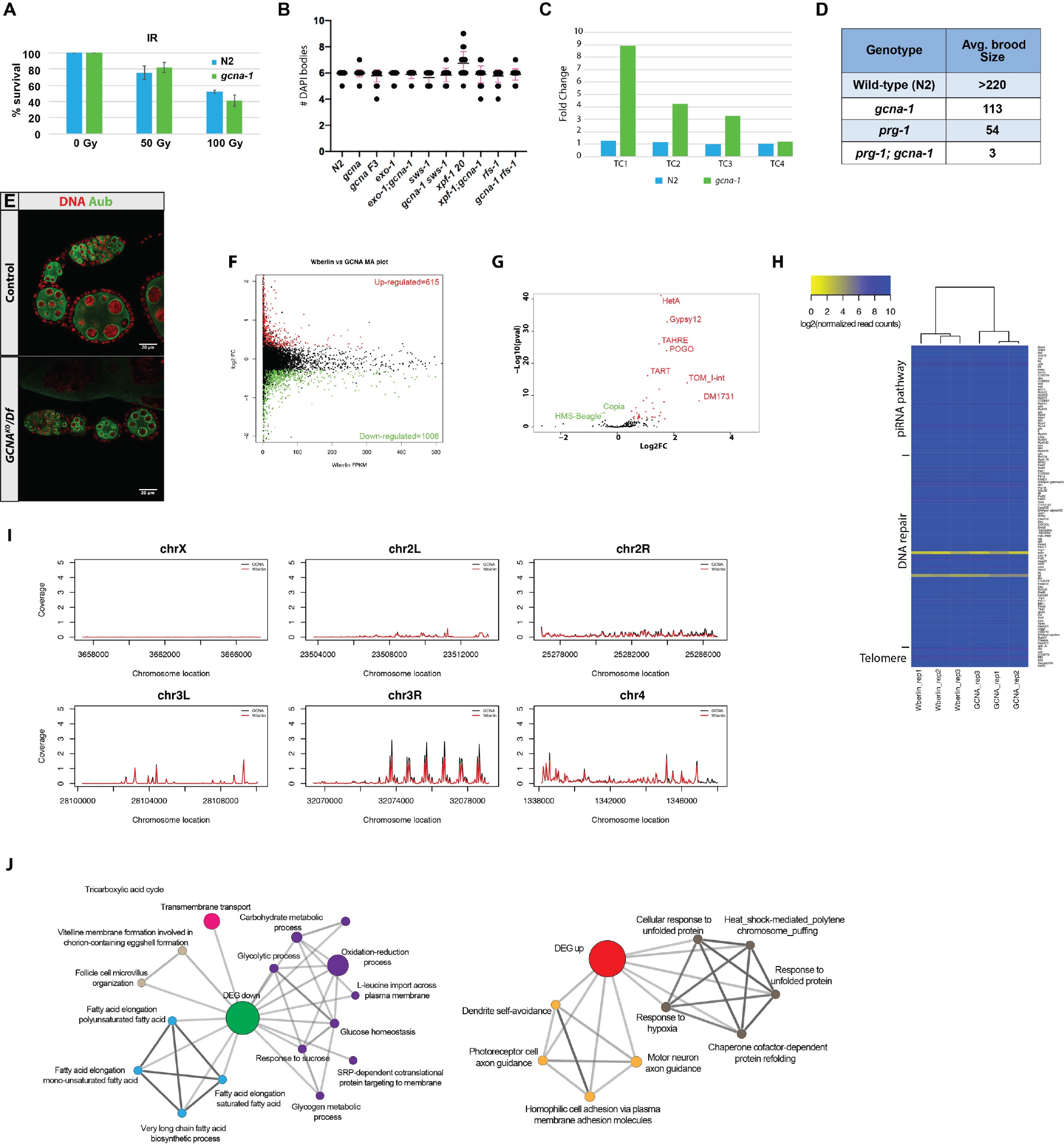
(related to Figure 3). *GCNA* mutants exhibit signs of replicative stress. **(A)** Percent survival of worms treated with the indicated dose of IR. **(B)** Quantification of DAPI bodies in the indicated genomic backgrounds. **(C)** Quantification of TE expression in *C. elegans* N2 control and *gcna-1* mutants. **(D)** Comparison of brood sizes of N2 (n=10), *gcna-1* (n=10), *prg-1* (n=10) and *prg-1; gcna-1* (N=20) mutant worms shifted from 20°C to 25°C as young L4 larvae. **(E)** Control and *GCNA*^*KO*^/*DF* ovaries stained for DNA (blue) and Aubergine (Aub) (Red). MA plot comparing gene expression in *GCNA*^*KO*^ and *w*^*Berlin*^ ovaries. **(G)** Volcano plot comparing TE expression in a *GCNA*^*KO*^ and *w*^*Berlin*^ ovaries. **(H)** Heat map comparing expression of 127 piRNA pathway, DNA repair and telomere maintenance genes. **(I)** Telomere visualization for all *Drosophila* chromosomes using average read density from *GCNA*^*KO*^ (black) and *w*^*Berlin*^ (red) samples. **(J)** GO analysis of the genes that exhibit down-regulation or up-regulation in a *GCNA*^*KO*^ background.

**Figure S4.**
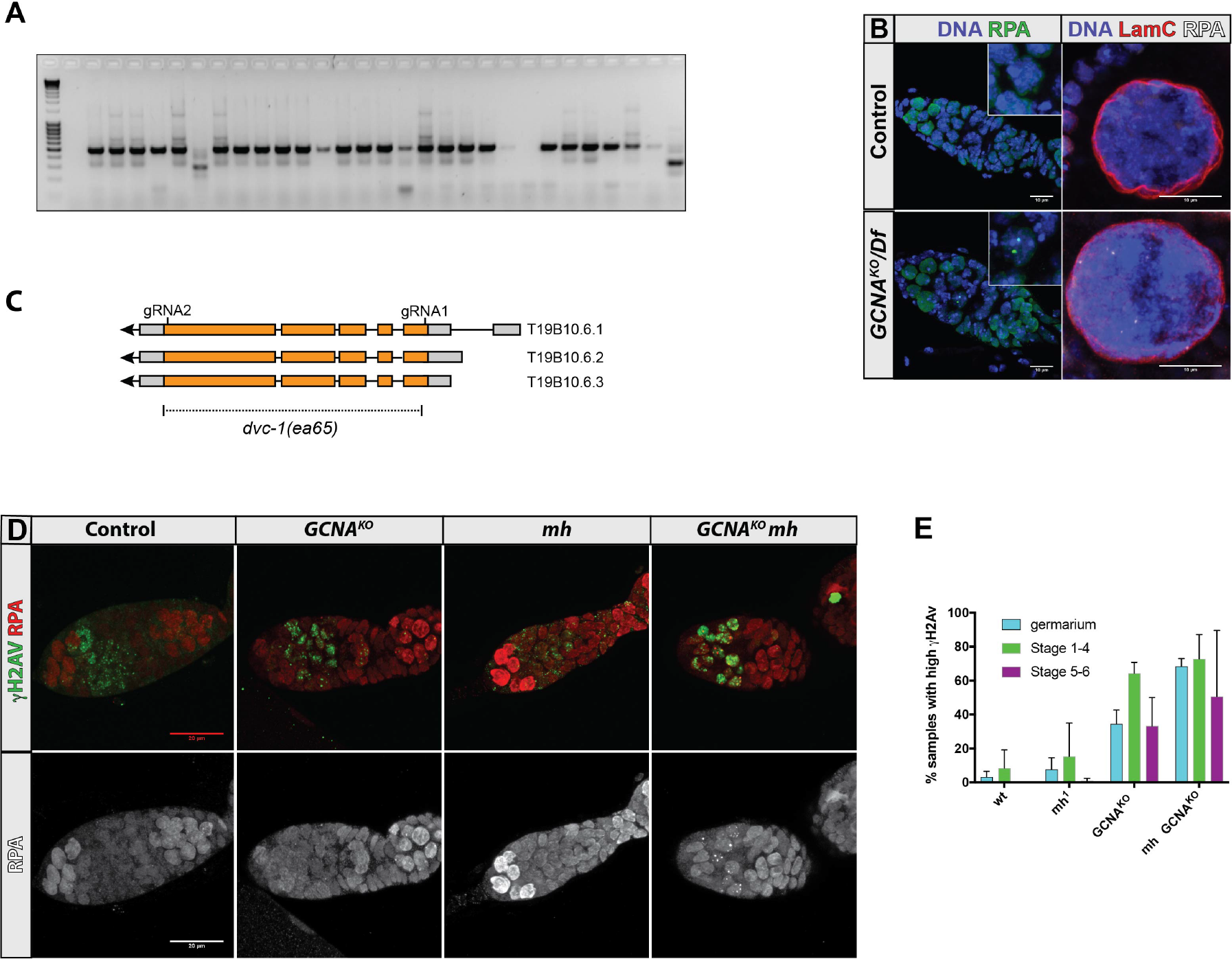
(related to Figure 3). *GCNA* mutants exhibit signs of replicative stress. **(A)** Representative gel showing PCR analysis of the *dog-1* assay. **(B)** Control and *GCNA*^*KO*^/*Df* germaria (left) and egg chamber nuclei (right) stained for DNA, RPA and Lamin C as indicated. **(C)** Structure of the *C. elegans dvc-1* gene locus and the *dvc-1(ea65)* allele. **(D)** Control, *GCNA*, *mh*, and *GCNA mh* mutants stained for RPA (red) and γH2Av (green). **(E)** Percent of *Drosophila* control, *GCNA*, *mh*, and *GCNA mh* mutant germline cysts at the indicated stages with high levels of nuclear γH2Av.

**Figure S5.**
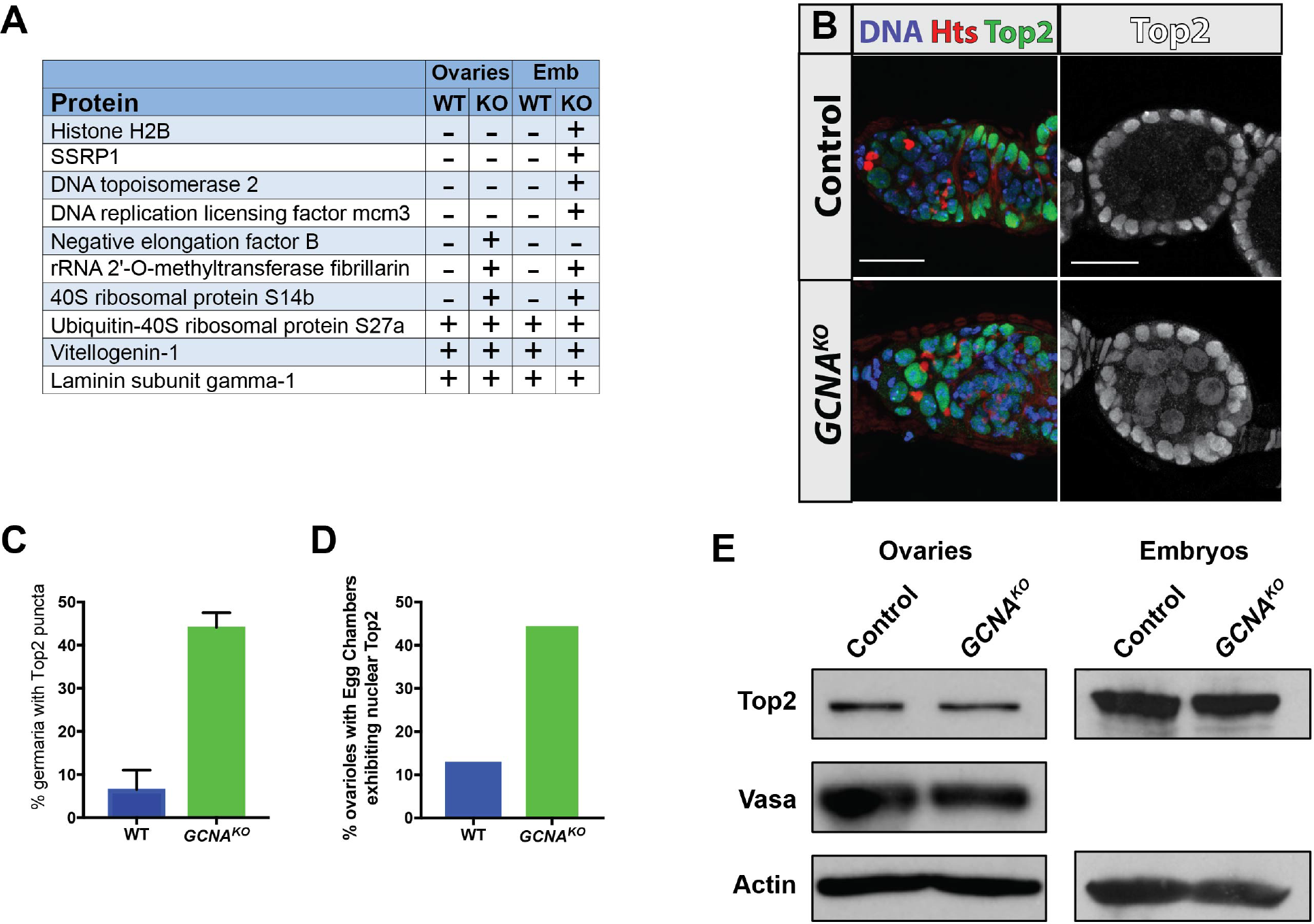
(related to Figure 4). GCNA mutants exhibit increased DPC levels. **(A)** Specific proteins were present in DPCs from wild-type and *GCNA*^*KO*^ *Drosophila* ovaries and embryos. **(B)** Control and *GCNA*^*KO*^ mutant ovaries stained for DNA (blue), Hts (red) and Top2 (green) **(C)** Quantification of germaria that contain germ cell nuclei with Top2 punctae. **(D)** Percent of ovarioles that contain egg chambers with clear germ cell nuclear Top2 localization. **(E)** Western blots of control and *GCNA*^*KO*^ mutant ovarian and embryo extracts probed for Top2, Vasa and Actin.

**Figure S6.**
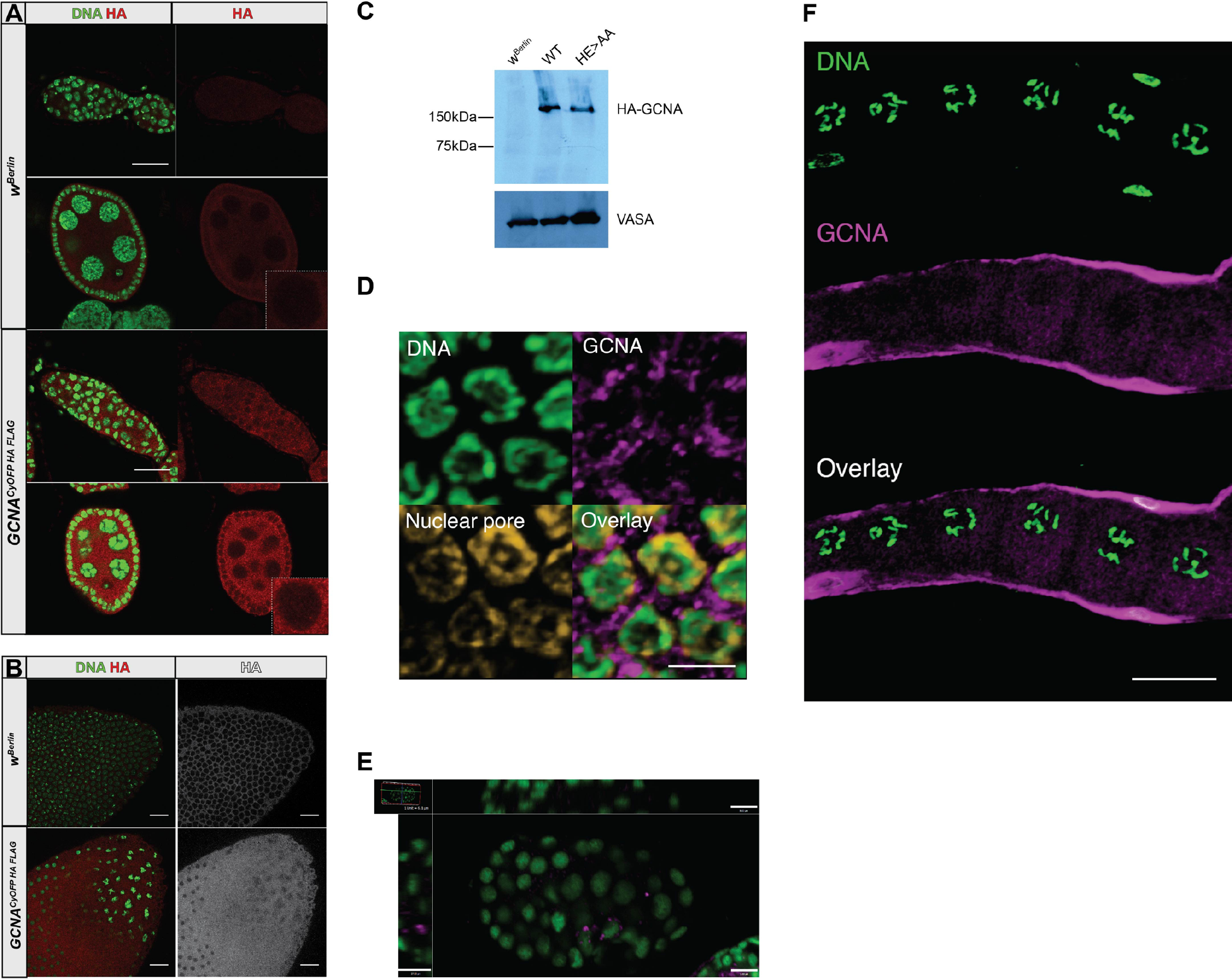
(related to Figure 5) GCNA expression in flies and worms. **(A)** Control *w*^*Berlin*^ and *GCNA*^*CyOFP HA FLAG*^ *Drosophila* ovaries stained for HA (red) and DNA (green). **(B)** Control *w*^*Berlin*^ and *GCNA*^*CyOFP HA FLAG*^ *Drosophila* embryos stained for HA (red) and DNA (green). **(C)** Western blot comparing expression of the HA-tagged *UAS GCNA*^*WT*^ and *GCNA*^*HE>AA*^ transgenes driven by *nanos* (*nos*)-*gal4* **(D)** Germ line squashes of *gcna-1(ea67* [*ollas::gcna-1*]) animals stained for DNA (DAPI, green), GCNA (anti-OLLAS, magenta), and nuclear pores (mAb414, yellow). Faint nucleoplasmic staining of GCNA-1 can be observed. **(E)** Primordial germ cell expression of GCNA-1 is seen in *gcna-1(ea67* [*ollas::gcna-1*]) embryos stained for DNA (green) and GCNA (anti-OLLAS, magenta). Shown are single plane image from a Z-stack. Scale 5µm. **(F)** *gcna-1(ea67* [*ollas::gcna-1*]) germ lines were stained for DNA (green) and GCNA (anti-OLLAS, magenta). Granular cytoplasmic staining is readily observed in the diplotene nuclei. Scale 20µm.

